# Interactome mapping reveals a role for LRP10 in autophagy and NDFIP1-mediated α-synuclein secretion

**DOI:** 10.1101/2023.11.28.569009

**Authors:** Ana Carreras Mascaro, Federico Ferraro, Valerie Boumeester, Guido Breedveld, Dick H.W. Dekkers, Leonie J.M. Vergouw, Frank Jan de Jong, Jeroen A. A. Demmers, Vincenzo Bonifati, Wim Mandemakers

## Abstract

Variants in the *LRP10* gene have been found in a spectrum of neurodegenerative disorders, including Lewy body diseases (LBDs). In brains of LBD patients, LRP10 is found in neuronal α-synuclein-containing Lewy bodies, astrocytes, and vasculature, but not in inclusion-free neurons. Furthermore, recent work suggests that LRP10 is involved in α-synuclein processing and transmission, which is disrupted by the LBD-associated *LRP10*:c.1424+5G>A variant (LRP10-Splice). In spite of the cumulating genetic and functional evidence for a role of LRP10 in neurodegenerative disorders, our knowledge about the biological processes in which LRP10 is involved is incomplete. In this work, we provide a list of LRP10 interactors identified via LRP10 co-immunoprecipitation and mass spectrometry in LRP10-overexpressing cells and induced pluripotent stem cells (iPSC)-derived astrocytes. In addition to interactors and biological processes previously associated with LRP10, we identified novel interactors and pathways that may provide new insights into LRP10 function. Based on these findings, we focused on the involvement of LRP10 in the autophagy and unconventional secretion pathways via its interaction with the autophagy receptor SQSTM1/p62 and the ubiquitin-proteasome adaptor protein NDFIP1, respectively. We demonstrate that changes in LRP10 levels, either via knock-out or overexpression, affect p62 levels and autophagy in HuTu-80 cells and iPSC-derived astrocytes. Furthermore, we found that both LRP10 and NDFIP1 stimulate α-synuclein secretion and synergistically affect intracellular α-synuclein levels. Next, we studied the LRP10 interactome and related biological processes in iPSC-derived astrocytes carrying the LRP10-Splice variant. Although various interactors and biological processes were shared between wild-type LRP10 (LRP10-WT) and LRP10-Splice, others were only found in either LRP10-WT or LRP10-Splice. Interestingly, we found that LRP10-Splice responded differently to autophagy-modulating drugs in comparison to LRP10-WT. Furthermore, we show that LRP10-Splice interferes with the LRP10-WT:NDFIP1 interaction and NDFIP1-mediated α-synuclein secretion. Finally, we investigated the interactome of a secreted LRP10 species only found in conditioned media from LRP10-Splice carrier cells, and identify biological processes that might be impacted by the secreted LRP10-Splice specific protein. In summary, this study enhances our understanding of LRP10 biology, describes LRP10 functions in autophagy and NDFIP1-mediated α-synuclein secretion, and reveals potentially interesting differences between LRP10-WT and LRP10-Splice carrier cells that might be relevant to better understand the role of LRP10 in LBDs pathogenesis.

## Introduction

The low-density lipoprotein receptor-related protein 10 (LRP10) is a single-pass transmembrane protein, and a member of the low-density lipoprotein receptor family (Pohlkamp et al., 2017). Rare variants in *LRP10* have been identified in patients diagnosed with autosomal dominant familial forms of Parkinson’s disease (PD), Parkinson’s disease dementia (PDD), and dementia with Lewy bodies (DLB) (Quadri et al., 2018). These diseases are referred to as Lewy body diseases (LBDs), and they share pathological hallmarks that include degeneration of dopaminergic neurons in the substantia nigra pars compacta and other brain areas, and the presence of intra-neuronal α-synuclein-containing aggregates called Lewy bodies (Attems et al., 2021). After the initial report (Quadri et al., 2018), additional, rare *LRP10* variants have been described in LBD patients (Chen et al., 2019; Gagliardi et al., 2021; Li et al., 2021; Liao et al., 2021; Manini et al., 2021; Neri et al., 2021; Periñán et al., 2020; Song et al., 2023; Tan et al., 2019; Vergouw et al., 2019; Zhao et al., 2021a; Zhao et al., 2020) and in a spectrum of neurodegenerative disorders including Alzheimer’s disease (AD) (Vergouw et al., 2020a), frontotemporal dementia (Daida et al., 2019), progressive supranuclear palsy (Vergouw et al., 2020b), and amyotrophic lateral sclerosis (Ni et al., 2021). In addition to these genetic findings, functional evidence associating LRP10 in processes relevant for neurodegenerative disorders has also been reported. Previous research has implicated this protein in the intracellular vesicle transport pathway given its localization and binding to clathrin adaptors (Boucher et al., 2008; Brodeur et al., 2009; Doray et al., 2008; Quadri et al., 2018; Steinberg et al., 2013). LRP10 has also been shown to colocalize with the PD-associated retromer protein VPS35 (Grochowska et al., 2021; Quadri et al., 2018), and loss of VPS35 expression results in loss of LRP10 localization at the plasma membrane and accumulation in lysosomes (Daly et al., 2023; Steinberg et al., 2013). Additionally, LRP10 has been described as a receptor for the amyloid precursor protein (Brodeur et al., 2012) and Apolipoprotein E (ApoE)-carrying lipoproteins (Sugiyama et al., 2000), and as an interactor of the AD-associated protein SORL1 (Grochowska et al., 2021). Moreover, transcriptomic studies have nominated LRP10 as a driver of a subtype of AD (Neff et al., 2021), and as a key regulator of cognitive dysfunction observed in AD patients across sex differences and ApoE-genotypes (Guo et al., 2023). Interestingly, a mouse model of PD was shown to present up-regulation of the ortholog *Lrp10* in α-synuclein inclusion-bearing neurons (Goralski et al., 2023). This is in line with the results of our neuropathological study that identified LRP10 immunoreactivity in 86% of mature Lewy bodies in the substantia nigra not only of *LRP10*-variant carriers but also of idiopathic PD and DLB patients (Grochowska et al., 2021). This finding is particularly intriguing, considering that in brains of control subjects LRP10 is expressed in non-neuronal cells, with the highest levels in astrocytes and neurovasculature, but is undetectable in neurons (Grochowska et al., 2021). Interestingly, we found that LRP10 is secreted via extracellular vesicles (EVs) through an autophagy-sensitive process, and autophagy blockade leads to the formation of aberrant LRP10-positive structures in the neurons of human midbrain-like organoids (Carreras Mascaro et al., 2023). Moreover, LRP10 was found to regulate α-synuclein intracellular levels and secretion and the LBD-associated *LRP10:*c.1424+5G>A variant had a dominant negative effect on this regulation (Carreras Mascaro et al., 2023). In summary, multiple lines of evidence support a role for LRP10 in neurodegeneration. However, our knowledge about LRP10 function in physiological and pathological conditions remains limited.

In this study, we performed a series of LRP10 co-immunoprecipitations followed by mass spectrometry (MS) in LRP10-overexpressing HEK-293T cells and induced pluripotent stem cells (iPSC)-derived astrocytes to identify LRP10 interacting proteins. We used this LRP10 interactome to nominate significant LRP10-associated biological processes, validate selected interactors, and explore the role of LRP10 and validated interactors in the associated biological processes. We identified the autophagy receptor SQSTM1/p62 and the ubiquitin-proteasome pathway adaptor NDFIP1 as novel interactors of LRP10. Moreover, we show that LRP10 has a modulating role in autophagic function and we provide new clues on LRP10 activity as a regulator of α-synuclein levels and secretion via its interaction with NDFIP1. Finally, we analyzed the LRP10 interactome and related biological processes in an iPSC-derived astrocyte line from a donor carrying the previously described LBD-associated variant *LRP10*:c.1424+5G>A (LRP10-Splice) (Quadri et al., 2018), which aberrantly secretes LRP10 as a high molecular weight species in an EVs-independent manner (Carreras Mascaro et al., 2023). We demonstrate that LRP10-Splice interferes with the autophagy pathway, the LRP10:NDFIP1 interaction, and α-synuclein secretion. Furthermore, we identified interactors of the secreted LRP10-Splice specific protein species, show that it contains wild-type LRP10 protein, and nominate a list of processes that might be impacted by this aberrant species. Taken together, the identification of the LRP10 interactome provides novel insights into biological processes associated with LRP10 function in health and in LBDs, pointing to new promising target pathways for disease-modifying interventions.

## Materials and Methods

### Cloning

To overexpress LRP10 for mass spectrometry experiments, the previously described pLVX-EF1α-LRP10-IRES-NeoR plasmid was used (Carreras Mascaro et al., 2023). Briefly, LRP10 sequence-verified cDNA was subcloned from the pcDNA™3.1-LRP10-V5-His-TOPO® plasmid (Quadri et al., 2018) into the pLVX-EF1α-IRES-mCherry plasmid (Takara Bio, 631987) via Gibson Assembly® (NEB) according to the manufacturer’s specifications. Next, mCherry was removed via MluI and MscI digestion, and replaced via Gibson Assembly® with the Neomycin resistance gene (NeoR) subcloned from the pcDNA™3·1-LRP10-V5-His-TOPO® plasmid. The negative control plasmid pLVX-EF1α-IRES-NeoR was generated by removing LRP10 from the pLVX-EF1α-LRP10-IRES-NeoR plasmid via restriction digestion, blunt end generation, and T4 ligation. To overexpress LRP10-Splice, the pLVX-EF1α-LRP10^splice^-IRES-NeoR plasmid described in Carreras Mascaro et al. (2023) was used. Briefly, the LRP10-Splice fragment was generated by PCR of specific regions of LRP10 from the pLVX-EF1α-LRP10-IRES-NeoR plasmid and placed into the same plasmid after restriction digestion and removal of LRP10-WT. To generate the LRP10-knock out (KO) lines, the pSpCas9-GFP-LRP10gRNA or pSpCas9-PuroR-LRP10gRNA plasmids were used (Carreras Mascaro et al., 2023; Grochowska et al., 2022; Grochowska et al., 2021). To generate the doxycycline-inducible LRP10 cell lines, the pCW57-LRP10-2A-PuroR was used. First, LRP10 was subcloned from the pLVX-EF1α-LRP10-IRES-NeoR plasmid and placed after the rtTA (reverse tetracycline-controlled transactivator)-Advanced promoter in the AgeI-digested pCW57-MCS1-2A-MCS2 (Addgene, 71782, from Adam Karpf) plasmid via Gibson Assembly®. For NDFIP1 overexpression experiments, the pCMV3-NDFIP1 cDNA expression plasmid was used (Sino Biological, HG14656-UT). The pCMVsport6-SNCA (Horizon, clone Id: 6147966) plasmid was used for α-synuclein overexpression experiments. All plasmids were verified by Sanger sequencing.

### HEK-293T and HuTu-80 cell lines cell culture

HEK-293T and HuTu-80 (CLS, 300218) cell lines were expanded in growth medium (DMEM or DMEM:F12, respectively; Thermo Fisher Scientific, 10% fetal bovine serum (FBS), 1% Penicillin-Streptomycin) at 37 °C/5% CO_2_. Cells were split every two-three days with Trypsin– EDTA (Gibco).

### HEK-293T and HuTu-80 cells transfection, stable lines generation, and treatments

For transfection, cells were seeded to a 60-70% confluency in tissue culture plates. Transfections were performed using the GeneJuice® transfection reagent (Merck) according to the manufacturer’s specifications, and the samples were further processed 48 h after transfection or when otherwise specified. To generate LRP10-expressing stable lines, LRP10-KO HuTu-80 cells were transfected with pCW57-LRP10-2A-PuroR, and kept in media with 1 µg/mL Puromycin (Invivogen, ant-pr-1). To induce LRP10 expression, cells were treated with 1 µg /mL doxycycline hyclate (Dox, Sigma-Aldrich, D9891) for 48 h. To assess autophagic function, cells were treated with Bafilomycin A1 (BafA1, 200 nM, Enzo, BML-CM110-0100) and/or Torin1 (200 nM, Invivogen, inh-tor1). All compounds were mixed with culture medium and placed onto plated cells for 4 h before extraction.

### LRP10 knock-out cell lines generation

LRP10-KO HEK-293T, HuTu-80, and NPCs were generated as described in (Grochowska et al., 2022) and (Carreras Mascaro et al., 2023), respectively. LRP10-KO HEK-293T and HuTu-80 cell lines were provided by Martyna M. Grochowska. Briefly, HEK-293T and HuTu-80 cells were transfected with the pSpCas9-GFP-LRP10gRNA plasmid using GeneJuice® reagent (Merck). After 48 h, cells were dissociated, and GFP-positive cells were sorted as single cells with the BD FACSAria™ III cell sorter. Recovered clones were expanded as independent clones and genotyped. To generate LRP10-KO NPCs, the Control 1 line was transfected with the pSpCas9-PuroR-LRP10gRNA plasmid using Lipofectamine™ Stem Transfection Reagent (Thermo Fisher Scientific). Twenty-four to 72 h after transfection they were selected with 0.5 µg/mL of Puromycin (Invivogen). Individual cells were allowed to recover into single colonies, which were independently expanded and genotyped. For the three cell types, we selected biallelically targeted clonal lines carrying a homozygous, single base-pair insertion, predicted to result in a premature stop codon in *LRP10* (p.Gly12Trp fs*18).

### Primary cell lines

Details regarding the control and patient clonal lines used in this study can be found in Table 1. The Control 1 cell line was obtained from the Gladstone Institutes, while the Control 2 line and the line derived from a DLB patient carrying the *LRP10*:c.1424+5G>A variant (Quadri et al., 2018) were obtained within study protocols approved by the medical ethical committee of Erasmus MC and conformed to the principles of the Declaration of Helsinki. The participating subjects provided written informed consent for the use of the material for research purposes.

**Table 1:**
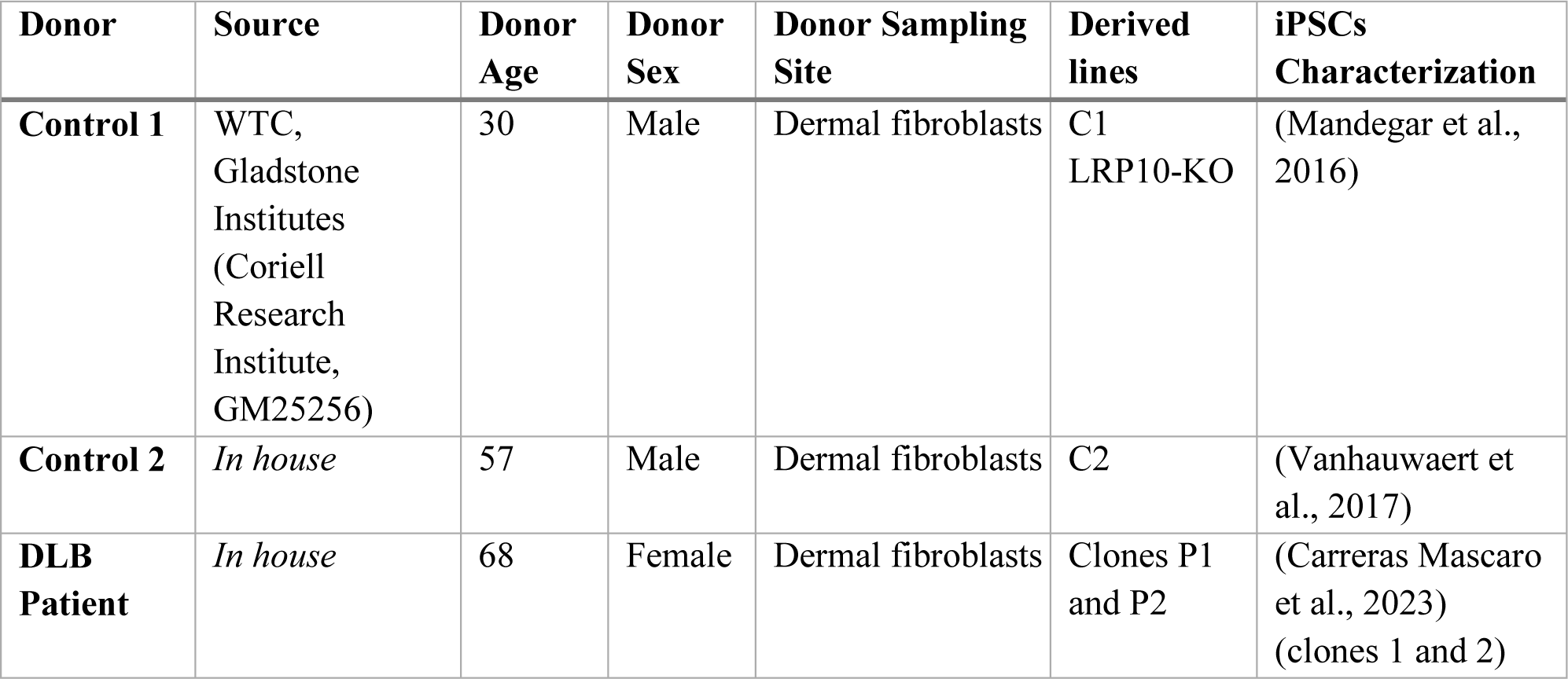
Details of the control and patient clonal lines included in this study.

| Donor | Source | Donor Age | Donor Sex | Donor Sampling Site | Derived lines | iPSCs Characterization |
| --- | --- | --- | --- | --- | --- | --- |
| Control 1 | WTC, Gladstone Institutes (Coriell Research Institute, GM25256) | 30 | Male | Dermal fibroblasts | C1<br>LRP10-KO | (Mandegar et al., 2016) |
| Control 2 | *In house* | 57 | Male | Dermal fibroblasts | C2 | (Vanhouwaert et al., 2017) |
| DLB Patient | *In house* | 68 | Female | Dermal fibroblasts | Clones P1 and P2 | (Carreras Mascaro et al., 2023)<br>(clones 1 and 2) |

### Generation of neural progenitor cells and astrocytes

Neural progenitor cells (NPCs) were provided by Martyna M. Grochowska and were generated as described in Grochowska et al. (Grochowska et al., 2021). Briefly, iPSCs were detached and cultured in a shaker as embryoid bodies. Neural identity was induced with the addition of small molecules and embryoid bodies were dissociated and plated on Matrigel® (Corning, 356238)-coated plates. NPCs were expanded in NPC medium: N2B27 medium (DMEM/F-12 – Neurobasal in 1:1 ratio, 1% B27 w/o Vitamin A, 0.5% N2, 1% Penicillin-Streptomycin, all from Thermo Fisher Scientific) supplemented with 3 μM CHIR, 200 μM ascorbic acid (AA, Sigma-Aldrich, A92902), and 0.5 μM Smoothened Agonist (SAG, Abcam, ab142160). Cells were kept on Matrigel-coated plates and split with accutase (Sigma-Aldrich) in a 1:10 ratio. NPCs were differentiated into astrocytes according to Carreras Mascaro et al. (Carreras Mascaro et al., 2023) Briefly, 75.000 NPCs/cm^2^ were seeded on Matrigel® (Corning)-coated plates with N2B27 medium supplemented with 10 ng/mL FGF-basic (PeproTech, 100-18B) and 10 ng/mL EGF (PeproTech, AF-100-15). After two days, cells were expanded in N2 medium (Advanced DMEM/F-12, 4% FBS, 1% N2, 1% Penicillin-Streptomycin) supplemented with 10 ng/mL CNTF (PeproTech, 450-13), which was switched to 10 ng/mL EGF after 12 days. Cells were kept in medium supplemented with EGF for 2-3 months. Three weeks before terminating cultures, cells were kept in medium supplemented with CNTF. One week before terminating cultures, cells were treated with 100 μM dbcAMP in N2 medium. Cells were cultured for at least 3 months. Astrocyte cultures were split with accutase (Sigma-Aldrich) at a 1:2-1:5 ratio when maximal density was reached.

### Protein lysates extraction

Cultured cells were washed with PBS and lysed with protein lysis buffer (100 mM NaCl, 1.0% Nonidet P-40 Substitute [Sigma, 74385], 50 mM Tris-Cl, pH 7.4) supplemented with protease inhibitors Complete® (Merck) and Pefabloc® SC (Merck). Lysates were snap-frozen, thawed on ice, and cleared by centrifugation at 10,000 × g for 10 min at 4 °C.

### EV-free media fraction isolation

iPSC-derived astrocytes were grown in exosome-free N2 medium (FBS was replaced by One Shot™ Fetal Bovine Serum, Exosome-Depleted, Gibco). Twenty milliliters of media were centrifuged at 2,000 × g for 10 min at 18 °C, and the supernatant was concentrated with Amicon Ultra-15 filters (Millipore, UFC901024) to a final volume of 10 mL. The concentrate was collected and centrifuged at 16,000 × g for 20 min at 4 °C. Next, the samples were filtered with a membrane pore size of 0.2 μm and centrifuged at 120,000 × g for 4 h at 4 °C in polycarbonate bottle assemblies (Beckman coulter, 355603) in a 70.1 Ti fixed angle rotor (Beckman coulter, 342184) in an Optima XE-90 ultracentrifuge (Beckman coulter). The final supernatant, or EV-free media fraction, was collected and further processed.

### Co-immunoprecipitation

The protein concentration of the cell lysates was determined with the Pierce™ BCA Protein Assay Kit (Thermo Fisher Scientific). For co-immunoprecipitation, Pierce™ Protein G Magnetic Beads (20 μL, Thermo Fisher Scientific Fisher) were washed three times with washing buffer (50 mM Tris-Cl (pH 7.4), 0.5 M NaCl, 0.05% v/v TWEEN® 20) and incubated with 2 μg of rabbit anti-LRP10 (Sino Biological, 13228-T16), 5 μg sheep anti-LRP10 (MRC PPU, DA058), or 5 μg sheep IgG isotype control (Invitrogen, 31243) antibody for 10 min at RT. Beads were washed three times with washing buffer and incubated with 400 µg protein lysates overnight at 4 °C. For conditioned media, 10 mL of media supernatant was concentrated 10 × with Amicon Ultra-15 filters (Millipore, UFC901024) to a final volume of 1 mL. Beads were washed three times with washing buffer and further processed for mass spectrometry or western blotting.

### Mass Spectrometry

The beads were first thoroughly washed in 50 mM ammonium bicarbonate (Sigma Aldrich). Proteins were then reduced in 50 mM Tris HCl (pH 8.2) / 2.4 mM sodium deoxycholate / 2.4 mM sodium N-lauryl sarcosine / 5 mM dithiothreitol (all from Sigma Aldrich). Sulfhydryl groups in cysteines were alkylated with 10 mM 2-chloroacetamide (Fluka, Honeywell Research Chemicals) to form carbamidomethyl derivatives. Samples were 1:1 diluted with 50 mM ammonium bicarbonate solution and overnight digestion was performed by adding 0.5 µg trypsin (TPCK Trypsin, Thermo Scientific) and 1 mM CaCl2 (Merck). Detergents were removed / precipitated in 0.5 % trifluoroacetic acid (Riedel-de Haën, Honeywell Research Chemicals) and the protein digests were washed twice with water saturated ethylacetate (Merck). Desalting of the proteolytic peptide sample was performed on a C18 solid phase extraction column (‘home-made ZipTip’ from a 15G punch from Empore™ octadecyl C18 47 mm extraction disks (3M, Bellefonte)) and after purification proteins were solubilized in 3 % acetonitrile and 0.5 % formic acid (both from Biosolve). Peptides were separated on a home-made 20 cm x 100 µm C18 column (BEH C18, 130 Å, 3.5 µm, Waters) after trapping on a nanoAcquity UPLC Symmetry C18 trapping column (Waters, 100 Å, 5 µm, 180 µm x 20 mm), using an EASY-nLC 1000 liquid-chromatograph. Subsequent mass spectrometry analyses were performed on a Thermo Scientific Orbitrap Fusion™ Lumos Tribrid™ mass spectrometer directly coupled to the EASY-nLC, as described (Sap et al., 2017).

For data dependent acquisition (DDA): All mass spectra were acquired in profile mode. The resolution in MS1 mode was set to 120,000 (AGC target: 4E5), the m/z range 350-1400. Fragmentation of precursors was performed in 2 s cycle time data-dependent mode by HCD with a precursor window of 1.6 m/z and a normalized collision energy of 30.0; MS2 spectra were recorded in the orbitrap at 30,000 resolution. Singly charged precursors were excluded from fragmentation and the dynamic exclusion was set to 60 seconds.

For data independent acquisition (DIA): all spectra were recorded at a resolution of 120,000 for full scans in the scan range from 350–1650 m/z. The maximum injection time was set to 50 ms (AGC target: 4E5). For MS2 acquisition, the mass range was set to 336–1391 m/z with dynamic isolation windows ranging from 7–82 m/z, with a window overlap of 1 m/z. The orbitrap resolution for MS2 scans was set to 30,000. The maximum injection time was at 54 ms (AGC target: 5E4; normalized AGC target: 100 %).

Raw mass spectrometry data were analyzed with the MaxQuant software suite (Cox et al., 2011) (version 2.1.3.0) as described previously (Schwertman et al., 2013) with the additional options ‘LFQ’ and ‘iBAQ’ selected. The A false discovery rate of 0.01 for proteins and peptides and a minimum peptide length of 7 amino acids were set. The Andromeda search engine was used to search the MS/MS spectra against the Uniprot database (taxonomy: Homo sapiens, release December 2022) concatenated with the reversed versions of all sequences. A maximum of two missed cleavages was allowed. The peptide tolerance was set to 10 ppm and the fragment ion tolerance was set to 0.6 Da for HCD spectra. The enzyme specificity was set to trypsin and cysteine carbamidomethylation was set as a fixed modification. Both the PSM and protein FDR were set to 0.01. In case the identified peptides of two proteins were the same or the identified peptides of one protein included all peptides of another protein, these proteins were combined by MaxQuant and reported as one protein group. Before further statistical analysis, the ‘proteingroups.txt’ table was filtered for contaminants and reverse hits (Supplementary Table 1).

DIA raw data files were analyzed with the Spectronaut Pulsar X software package (Biognosys, version 17.0.221202), using directDIA for DIA analysis including MaxLFQ as the LFQ method and Spectronaut IDPicker algorithm for protein inference. The Q-value cutoff at precursor and protein level was set to 0.01. All imputation of missing values was disabled. The logarithm (log 2) of the LFQ values were taken, resulting in a Gaussian distribution of the data (Supplementary Fig. 1). Zipped MaxQuant data have been deposited to the PRIDE repository with the data identifier PXD045871 and a summary is reported in Supplementary Table 1.

### Functional enrichment analysis

All data sets were analyzed in R v4.2.1.

Functional enrichment analysis was performed using the clusterProfiler v4.4.4 (Wu et al., 2021) and org.Hs.eg.db v3.15.0 R packages with the Gene Ontology Biological Process (GOBP) data set. All resulting p-values were corrected using the Benjamini-Hochberg (*BHp*-value) method.

Statistically significant GOBP terms were summarized with GOSemSim v2.24.0 (Yu, 2020; Yu et al., 2010) and rrvgp v1.8.1 (Sayols, 2023). Semantic similarity among the GOBP terms was measured using the graph-based method by Wang (Wang et al., 2007), then similar terms were summarized with a threshold of 0.9.

### Western blotting

Protein lysates were added 4 × sample buffer (8% SDS, 20% v/v glycerol, 0.002% bromophenol blue, 62.5 mM Tris-Cl, pH 6.8) supplemented with 400 mM dithiothreitol (DTT; except conditioned media samples, which were not supplemented with DTT) and incubated for 10 min at 95 °C. For co-immunoprecipitations, the beads were eluted with 30 μL of 1 × sample buffer (2% SDS, 5% v/v glycerol, 0.0005% bromophenol blue, 15.6 mM Tris-Cl (pH 6.8)) supplemented with 100 mM DTT for 10 min at 95 °C. Beads were magnetized and discarded. Proteins were separated on 4-15% Criterion TGX precast gels (Bio-Rad). Gels were transferred to nitrocellulose membranes using the Trans-Blot® Turbo™ Transfer System (Bio-Rad). Membranes were blocked using 5% non-fat dry milk (Blotto, Santa Cruz Biotechnology) in Tris-buffered saline (TBS), 0.1% v/v TWEEN® 20 (Merck) for 1 h at room temperature. Next, they were incubated in blocking buffer containing the diluted primary antibodies overnight at 4 °C. After washing in TBS, 0.1% v/v TWEEN® 20, blots were incubated for 1 h at room temperature with fluorescently conjugated secondary antibodies. Membranes were washed with TBS, 0.1% v/v TWEEN® 20 and imaged using the Odyssey CLx Imaging system (LI-COR Biosciences).

### Immunocytochemistry of 2D cultures (ICC)

Astrocytes were plated on glass coverslips, fixed with 4% PFA for 10 min, and washed three times with PBS. Next, cells were incubated in blocking buffer (50 mM Tris.HCl, 0.9% NaCl, 0.25% gelatin, 0.2% Triton™ X-100, pH 7.4) containing primary antibodies at 4 °C overnight. The coverslips were then washed by dipping them in a PBS, 0.05% TWEEN® 20 solution. Cells were incubated for 1 h with fluorescently conjugated secondary antibodies at room temperature. The coverslips were washed with PBS, 0.05% TWEEN® 20, and incubated for 3 min with 0.1% (w/v) Sudan Black B (Sigma) in 70% ethanol. After a final washing with PBS, 0.05% TWEEN® 20 cells were mounted with ProLong Gold with DAPI (Invitrogen).

### Primary and secondary antibodies

Primary antibodies used for western Blot (WB) and immunocytochemistry (ICC) are listed in Table 2.

**Table 2:** Primary antibodies used for immunocytochemistry (ICC) and western blotting (WB) experiments.

| Target protein | Host | Supplier | Catalogue number | WB dilution | ICC dilution |
| --- | --- | --- | --- | --- | --- |
| **CD63** | Mouse | Thermo Fisher Scientific | 10628D | - | 1:200 |
| **GFAP** | Rabbit | Sigma-Aldrich | AB5804 | - | 1:200 |
| **LC3B** | Rabbit | Novus | NB100-2220 | 1:1000 | 1:100 |
| **LRP10** | Rabbit | Sino Biological | 13228-T16 | 1:1000 | - |
| **LRP10** | Sheep | MRC PPU | DA058 | 1:1000 | 1:100 |
| **NDFIP1** | Rabbit | Sigma-Aldrich | HPA009682 | 1:1000 | 1:100 |
| **p62** | Mouse | BD Biosciences | 610833 | 1:1000 | - |
| **s100β** | Mouse | Sigma-Aldrich | S2532 | - | 1:200 |
| **vinculin** | Mouse | Santa Cruz Biotechnology | 2679S | 1:5000 | - |
| **α-Synuclein** | Mouse | BD Biosciences | 610787 | 1:1000 | - |

Secondary antibodies used for WB: Alexa Fluor Plus 680 or 800 donkey anti-mouse and donkey anti-rabbit (all from Thermo Fisher ScientificFisher, 1:1000), or Alexa Fluor® 790 AffiniPure donkey anti-sheep (Jackson immunoresearch, 1:1000).

Secondary antibodies used for immunocytochemistry: Alexa Fluor® 488 donkey anti-sheep, Alexa Fluor® 594 donkey anti-mouse, Alexa Fluor® 647 goat anti-rabbit (all from Jackson ImmunoResearch Laboratories).

### Microscopy images acquisition and image analysis

Stainings were imaged with the Leica SP5 AOBS confocal microscope. The following lasers were used: diode 405, OPSL 488, DPSS 561, and HeNe 633. Each image was detected on the spectral PMT detector with an HCX PL APO CS 40 ×/1.25 or HCX PL APO CS 63 ×/1.4 lens. Scanning of detailed images was done with a pixel size of 0.1 μm and with a scan size of 2048 × 2048 pixels at 400 or 600 Hz. For z-stack images, 0.35 μm steps in the z-direction were taken. The pinhole size was set to 1 airy unit (AU). Microscopy images were analysed using Fiji/ImageJ version 1.45b. The built-in tool coloc2 was used to determine the Manders’ overlap coefficient between LRP10 and LC3B after generating maximum projections and automatic thresholding of each channel.

### Statistical analysis

Statistical analyses of all experiments excluding functional enrichment analyses were performed using Prism 9 software (GraphPad). Students T-test was used for experiments with only two conditions. For experiments with two or more groups, One-way or Two-way ANOVA followed by the Dunnett’s, Tukey’s, or Sidak’s multiple comparisons tests were used as specified in the figure legends. The Grubb’s test was used to discard outliers. Significant P values of *P* ≤ 0.05 were reported. In the graphs, “*” represents a p-value of *P* ≤ 0.05, “**” represents *P* ≤ 0.01, “***” represents *P* ≤ 0.001, and “****” represents *P* ≤ 0.0001. The data are presented as means ± standard deviations (SD) and represent results from at least 3 biological replicates.

## Results

### LRP10 interactome in HEK-293T cells reveals links to LBD-relevant pathways

Despite the recent advancements in studying the role of LRP10 in neurodegeneration, understanding of LRP10 physiological function remains limited. To determine biological pathways where LRP10 plays a potentially relevant role, we mapped the LRP10 interaction network via mass spectrometry (MS) of proteins that co-immunoprecipitated with LRP10.

We performed LRP10 immunoprecipitations using whole-cell extracts from LRP10-overexpressing (LRP10-OE), and LRP10 knock-out (LRP10-KO) HEK-293T cells, and subsequently performed MS analyses. Proteins detected in LRP10-KO HEK-293T cells were excluded from the list of the putative LRP10-interacting proteins. We performed the Co-IP/MS experiments twice for both LRP10-OE and LRP10-KO HEK-293T cells.

After quality control and filtering the proteins identified in the negative control experiments, we identified a total of 583 unique proteins across the two experiments, with 172 identified in both (Supplementary Table 2A). It is noteworthy that LRP10 was only detected in LRP10-OE and not in LRP10-KO cells. The LRP10 interactome in HEK-293T cells was significantly enriched (*BHp*-value < 0.05) for 427 GOBP terms that were further summarized into 27 summary terms (Fig. 1A, Supplementary Table 3). These terms included diverse functions among which some of them are already known to play a role in the pathophysiology of LBDs, e.g. protein folding, proteasome processes, mitochondrial biology, energy metabolism, autophagy, and substantia nigra development.

**Figure 1:**
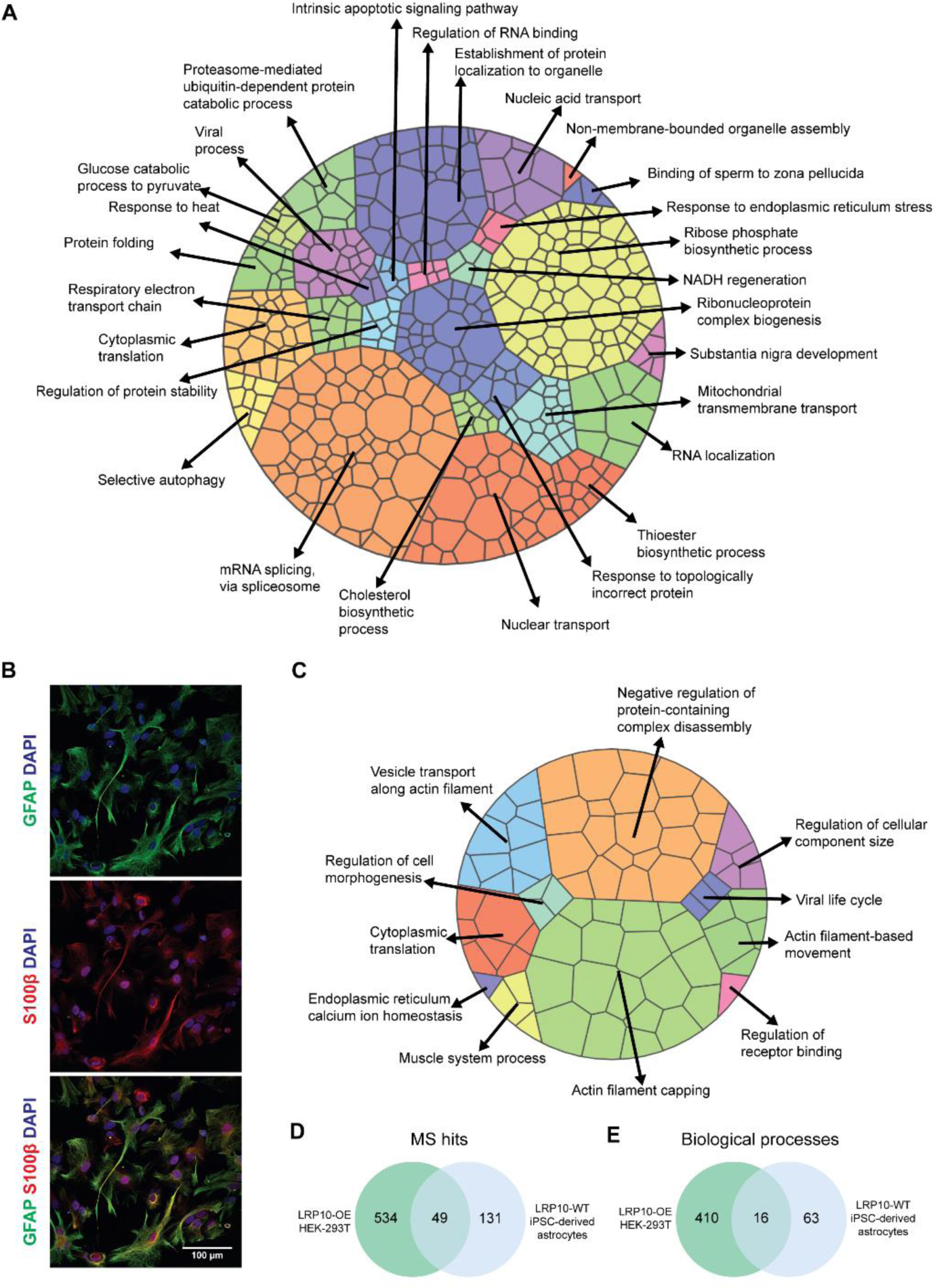
LRP10 interactome MS in LRP10-OE HEK-293T cells and LRP10-WT iPSC-derived astrocytes. **A**. Voronoi plot showing the 427 statistically significant GOBP identified via the LRP10-OE HEK29T cells co-immunoprecipitations/MS experiments. Each tile represents a GOBP term with a size that is inversely proportional to the *BHp*-value. The tiles are grouped into the 27 summary terms and labeled with the corresponding GOBP term. **B**. ICC characterization of 4-months old astrocytes showing the astrocytic markers GFAP and S100β. **C**. Voronoi plot showing the 79 statistically significant GOBP enriched in LRP10-WT iPSC-derived astrocytes. The tiles are grouped into the 11 summary terms. **D**. Venn diagram showing the overlap between the proteins identified in LRP10-OE HEK-293T cells and LRP10-WT astrocytes **E**. Venn diagram showing the overlap between the significantly enriched biological processes in LRP10-OE HEK-293T cells and LRP10-WT astrocytes.

### Mapping the LRP10 interactome in iPSC-derived astrocytes from healthy controls

LRP10 presents the highest expression levels in astrocytes and neurovasculature in brains from healthy individuals (Barbar et al., 2020; Grochowska et al., 2021; La Manno et al., 2016; Zhang et al., 2016). To determine LRP10-interacting proteins in a more physiologically-relevant system, we generated iPSC-derived astrocytes following established protocols (Carreras Mascaro et al., 2023; Grochowska et al., 2021; Reinhardt et al., 2013). iPSC-derived astrocytes were characterized for the presence of the astrocytic markers GFAP and S100β after 4 months of differentiation via immunocytochemistry (Fig. 1B). Next, we performed LRP10 immunoprecipitations coupled with MS in whole-cell extracts from two control (LRP10-wild-type (WT)) and LRP10-KO iPSC-derived astrocytes (Table 1). After removing all the hits that were also present in LRP10-KO astrocytes, we identified 180 LRP10 interactors (Supplementary Table 2B). Of note, LRP10 was only detected in controls and not in the LRP10-KO sample. Astrocytic LRP10 interactors were significantly enriched (*BHp*-value < 0.05) for 79 GOBP terms belonging to 11 summarized terms (Fig. 1C), including cytoplasmic translation, actin filament capping, negative regulation of protein-containing complex disassembly, actin filament-based movement, vesicle transport along actin filament, regulation of cellular component size, regulation of cell morphogenesis, muscle system process, viral life cycle, regulation of receptor binding, and endoplasmic reticulum calcium ion homeostasis. Compared with the results of the LRP10-OE HEK-293T experiment, 49 proteins were identified across the two cellular systems (Fig 1D, Table 3). Sixteen biological processes were significantly enriched in both experiments, even though none of the summarized terms overlapped (Fig 1E, Table 3). These processes included cytoskeleton biology, translation, calcium homeostasis, and viral processes. Taken together, the LRP10 interactome is associated with a collection of LBDs relevant proteins and biological processes in two independent LRP10 expressing cell models.

**Table 3:** LRP10-interacting proteins and significant biological processes identified in whole-cell extracts from both LRP10-OE HEK-293T and LRP10-WT astrocytes.

| Proteins (49) | Biological processes (16) |
| --- | --- |
| **ABCF2 ARL8B ATP1A1 ATP2A2**<br>**CAND1 CANX CNP COPA DPM1**<br>**DYNC1H1 ESYT2 GJA1 HACD3**<br>**HNRNPF HNRNPU IPO5 IPO7 IPO9**<br>**ITCH ITM2B KIDINS220 KPNB1**<br>**LRP10 MARCKSL1 NDFIP1 PRKDC,**<br>**RPL7A RPL8 RPL10A RPL14 RPL15**<br>**RPN1 RPN2 RTCB SLC16A1 SLC25A1**<br>**SLC25A4 SLC25A11 SLC39A7 SRSF3**<br>**TECR TMED10 TMEM59 TUFM**<br>**UQCRC2 USMG5 VAPA WWP2**<br>**YWHAQ** | cellular component disassembly, cytoplasmic translation,<br>cytoskeleton-dependent intracellular transport,<br>endoplasmic reticulum calcium ion homeostasis,<br>establishment of organelle localization,<br>movement in host, ncRNA processing,<br>peptidyl-asparagine modification,<br>protein N-linked glycosylation via asparagine,<br>ribonucleoprotein complex biogenesis,<br>ribosomal large subunit biogenesis,<br>ribosome biogenesis, rRNA metabolic process,<br>rRNA processing, viral life cycle, viral process |

### LRP10 interacts with p62 and modulates autophagic function

MS analysis in HEK-293T cells revealed that LRP10 interacts with several proteins involved in the autophagy pathway (Fig. 1A, Supplementary Table 2A), including the autophagy receptor p62 (Hou et al., 2020; Ma et al., 2019). Dysregulation of cellular degradation pathways, such as the autophagy and ubiquitin-proteasome pathways, is a very well-described phenomenon in LBDs (Behl et al., 2022; Hou et al., 2020). Furthermore, LRP10 levels and secretion were found to be dependent on autophagic function (Carreras Mascaro et al., 2023). To explore the role of LRP10 in the autophagy pathway in more detail, we performed LRP10 co-immunoprecipitation in LRP10-OE HEK-293T cells. We observed the presence of p62 after LRP10 immunoprecipitation only in LRP10-OE cells and not in LRP10-KO cells (Fig. 2A). To validate the LRP10:p62 interaction in a more physiological context, we performed LRP10 co-immunoprecipitation in iPSC-derived astrocytes even though p62 was not among the LRP10 interactors detected by MS in these cells. We have previously shown that LRP10 protein levels are strongly regulated by the autophagy-lysosomal pathway in iPSC-derived astrocytes (Carreras Mascaro et al., 2023), and reasoned that the LRP10:p62 interaction might be undetectable with our experimental setup in astrocytes when autophagy is active under physiological conditions. To test this possibility, we treated 4 months-old control iPSC-derived astrocytes with Bafilomycin (BafA1), which blocks the acidification of lysosomes and autolysosomes, for 4 h in order to minimize the breakdown of LRP10 and p62-containing protein complexes and facilitate their interaction. We performed an LRP10 immunoprecipitation with protein extracts from these cells and we observed that p62 co-immunoprecipitated with LRP10 in control cells but not in LRP10-KO cells (Fig. 2B).

**Figure 2:**
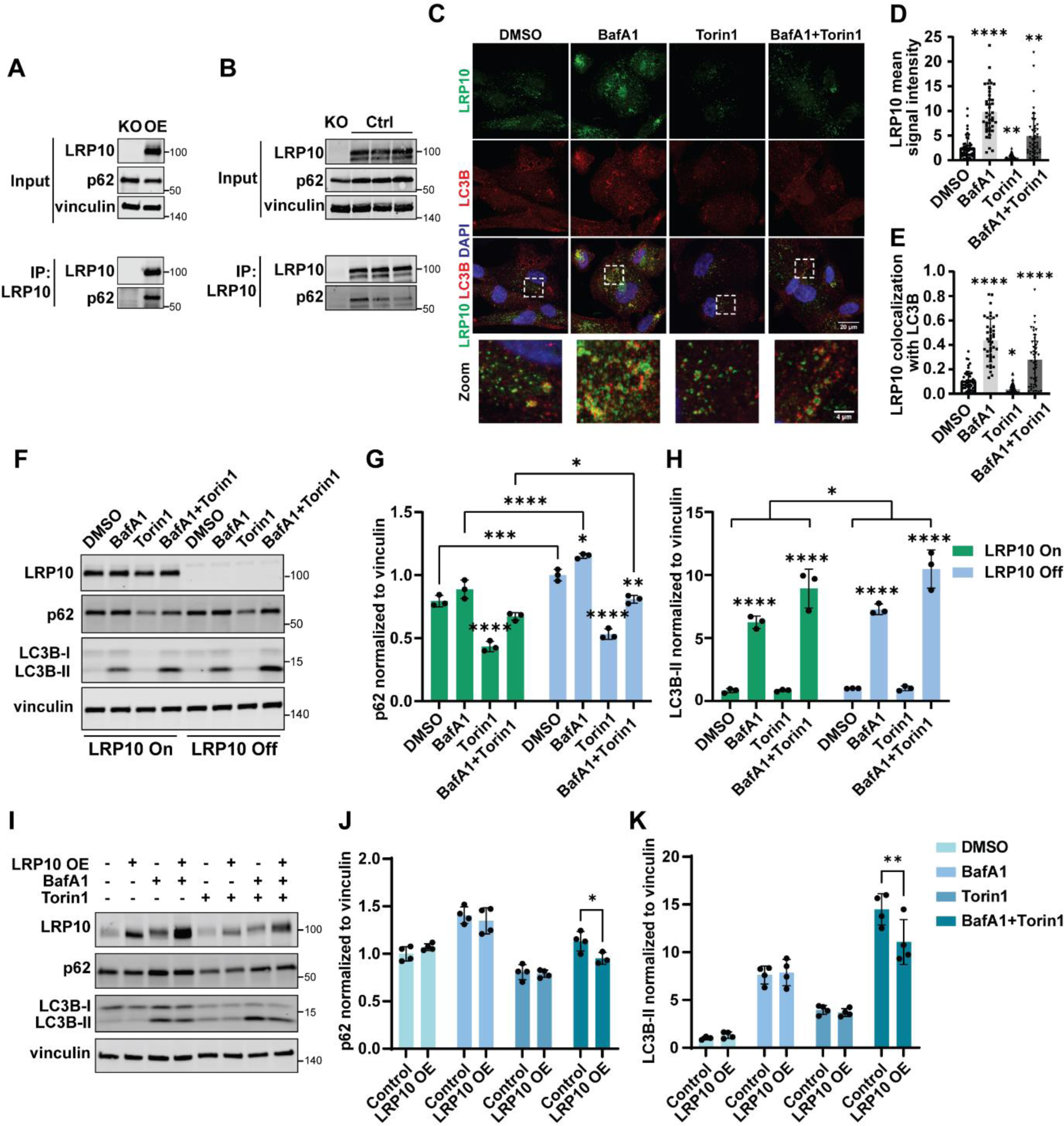
LRP10 levels influence autophagic function. **A**. Western blot showing LRP10 and endogenous p62 co-immunoprecipitation (IP) in LRP10-KO or LRP10-OE HEK-293T cells using an antibody directed against the C-terminal domain of LRP10 for immunoprecipitation. **B**. Western blot of endogenous LRP10 and endogenous p62 co-immunoprecipitation in LRP10-KO or LRP10-WT (Ctrl) 4 months-old iPSC-derived astrocytes after a 4 h BafA1 treatment. **C**. Immunocytochemistry of LRP10 and LC3B in 4 months-old astrocytes treated with DMSO, BafA1, Torin1, or BafA1+Torin1 for 4 h. **D**. Quantification of LRP10 mean signal intensity per cell from **C**. n > 40 cells per condition. **E**. Quantification of LRP10 colocalization (Mander’s overlap coefficient) with LC3B per cell from **C**. n > 40 cells per condition. **F**. Representative western blot of HuTu-80 stable cell lines expressing LRP10 (LRP10-On) or not (LRP10 Off) after 4 h treatments with DMSO, BafA1, Torin1, or a combination of BafA1+Torin1. **G**. Western blot quantification of p62 from F. n = 3 biological replicates. **H**. Western blot quantification of LC3B-II from F. n = 3 biological replicates. **I**. Representative western blot of control iPSC-derived astrocytes overexpressing LRP10 or not after 4 h treatments with DMSO, BafA1, Torin1, or BafA1+Torin1. **J**. Western blot quantification of p62 from I. n = 4 biological replicates. **K**. Western blot quantification of LC3B-II from I. n = 4 biological replicates. All data are expressed as mean ± SD with individual data points shown. Data from D and E were analyzed One-way ANOVA with Dunnett’s multiple comparisons test. Data from G, H, J, and K were analyzed by Two-way ANOVA with Tukey’s multiple comparisons test. *P ≤ 0.05, **P ≤ 0.01, ***P ≤ 0.001, ****P ≤ 0.0001.

Furthermore, we investigated the effects of the autophagy-modifying drugs BafA1 and Torin1 (a potent and selective ATP-competitive mTOR inhibitor) on LRP10 expression and localization in astrocytes via immunocytochemistry (Fig. 2C). In DMSO-treated cells, LRP10 localized to vesicular structures that partially overlapped with the autophagic marker LC3B (Fig. 2C, 2E). After BafA1 treatment, LRP10-positive vesicles accumulated in clusters and appeared enlarged (Fig. 2C). Image quantification revealed that BafA1 led to a significant 4-fold increase in both LRP10 signal intensity and LRP10 colocalization with LC3B (Fig. 2D, 2E), which suggests that the degradation of LRP10 and LC3B was inhibited and these proteins were accumulating in autophagosomes. In contrast, Torin1 treatment led to the formation of small vesicular structures with a 4-fold decrease in LRP10 signal and a 3-fold decrease in LRP10 colocalization with LC3B (Fig. 2C-E), indicating that LRP10 is degraded faster after autophagy induction. Simultaneous treatment with BafA1 and Torin1 lead to an intermediate state with enlarged and intense LRP10 structures (3-fold increase compared to DMSO) that were surrounded by LC3B-positive vesicles (Fig. 2C, 2D). Colocalization of LRP10 with LC3B was 3-fold higher in the BafA1+Torin1 condition compared to DMSO (Fig. 2E). Altogether, these data indicate that LRP10 interacts with p62 and is degraded via the autophagy pathway in iPSC-derived astrocytes.

To investigate whether LRP10 itself might modulate autophagy, we generated a Doxycycline (Dox)-inducible LRP10 expression stable cell line in LRP10-KO HuTu-80 cells and induced LRP10 expression for 48 hours until reaching expression levels comparable to endogenous LRP10 expression in HuTu-80-WT cells (Supplementary Fig. 2). To test autophagic function, the cells were treated for 4 hours with BafA1 and Torin1 and the levels of the autophagy indicators p62 and LC3B-II were inspected via western blot (Fig. 2F). In line with previous publications for other cell types, BafA1 induced an increase in p62 and LC3B-II levels, Torin1 induced a decrease in p62 and LC3B-I levels, and BafA1 combined with Torin1 induced a decrease in p62 levels but an increase in LC3B-II levels (Klionsky et al., 2021) (Fig. 2F). Interestingly, we observed significantly higher levels of p62 in DMSO, BafA1, and BafA1 and Torin1-treated LRP10-KO (LRP10 Off) cells in comparison to LRP10-expressing cells (LRP10 On) (Fig. 2F, 2G). Additionally, LC3B-II levels were significantly higher in LRP10 Off cells compared to LRP10-expressing cells (Fig. 2F, 2H). These results show that changes in LRP10 levels affect autophagic function, suggesting that LRP10 modulates autophagy.

To provide more evidence that LRP10 plays a relevant role in autophagic function, we overexpressed LRP10 in iPSC-derived astrocytes and treated them for 4 h with the autophagy-modifying compounds. To determine the effect of LRP10 overexpression, p62 and LC3B-II levels were assessed via western blotting (Fig. 2I). We observed significantly reduced p62 and LC3B-II levels in LRP10-overexpressing cells after the combined treatment with BafA1 and Torin1 (Fig. 2J, 2K). These results suggest that LRP10 can also modulate the autophagy pathway in iPSC-derived astrocytes, possibly via its interaction with p62. In summary, the observations showing that the lack of LRP10 in HuTu-80 cells leads to increased levels of p62 and LC3B-II, and LRP10 overexpression in iPSC-derived astrocytes induces a decrease in p62 and LC3B-II in BafA1+Torin1-treated cells suggest that LRP10 expression has a stimulatory effect on the autophagy pathway.

### LRP10 interacts with NDFIP1 to regulate α-synuclein levels and secretion

Nedd4 family-interacting protein 1 (NDFIP1) was detected as a LRP10 interactor in both HEK-293T cells and iPSC-derived astrocytes (Fig. 1, Supplementary Table 2D). NDFIP1 is a crucial molecule in the ubiquitin-proteasome pathway, it has been described to play a role in PD pathology (Howitt et al., 2014), and to be necessary for α-synuclein degradation (Borland et al., 2022; Liu et al., 2020). Importantly, α-synuclein is central in LBDs pathogenesis (Attems et al., 2021; Burré et al., 2018) and LRP10 has recently been shown to regulate α-synuclein intracellular levels and secretion (Carreras Mascaro et al., 2023). Based on this evidence, the interaction between LRP10 and NDFIP1 was further investigated.

First, to validate the physical interaction between LRP10 and NDFIP1 as identified via MS, we performed LRP10 immunoprecipitation in LRP10-KO and LRP10-OE HEK-293T cells and we observed NDFIP1 co-immunoprecipitating with LRP10 only in LRP10-OE cells (Fig. 3A).

**Figure 3:**
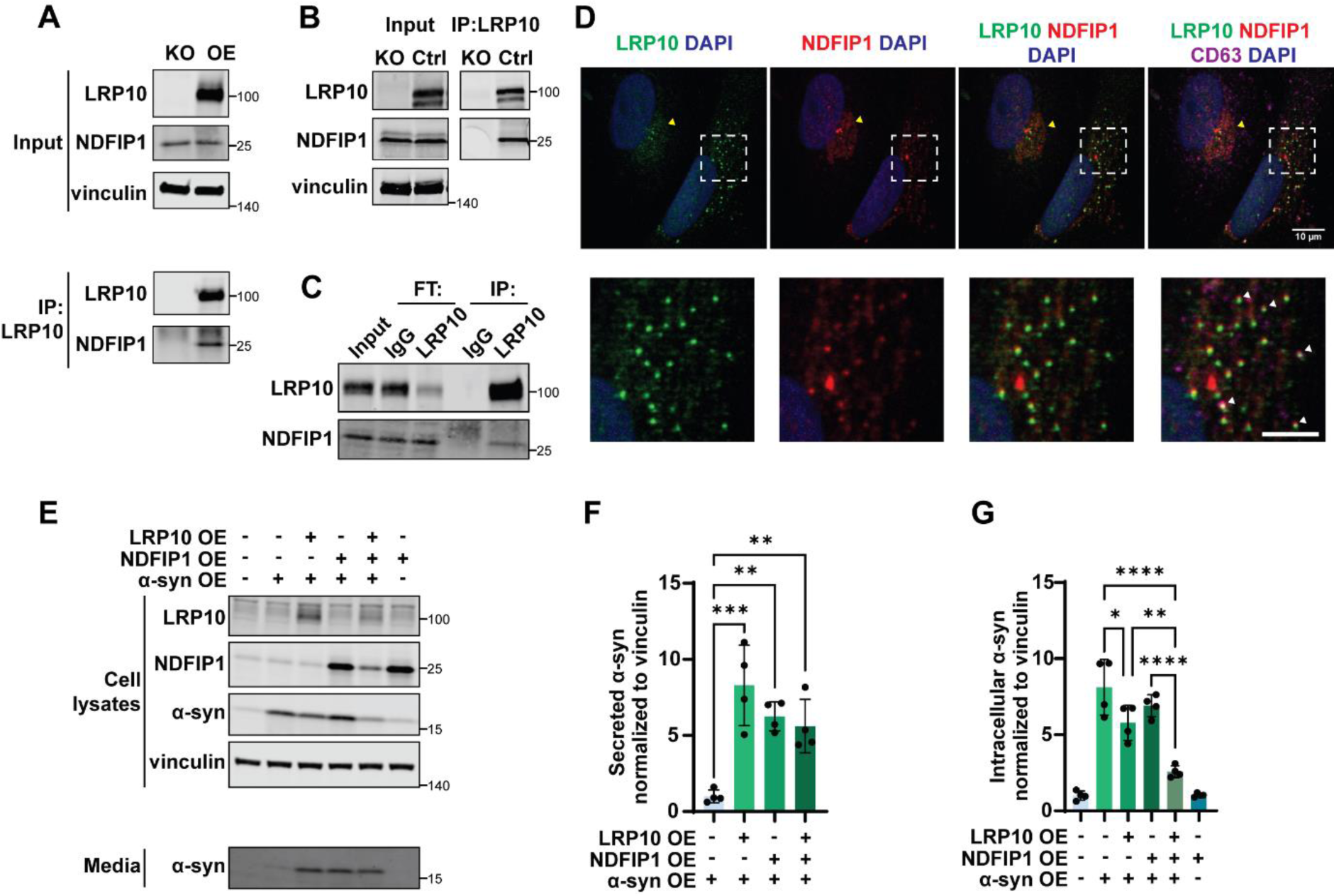
LRP10 interaction with NDFIP1. **A**. Western blot of endogenous NDFIP1 and LRP10 co-immunoprecipitation in LRP10-KO and LRP10-OE cells. An antibody directed against the C-terminal domain of LRP10 was used for immunoprecipitation. **B**. Endogenous LRP10 and NDFIP1 co-immunoprecipitation in 4 months old LRP10-KO and control iPSC-derived astrocytes. An antibody directed against the C-terminal domain of LRP10 was used for immunoprecipitation. **C**. Endogenous LRP10 and NDFIP1 co-immunoprecipitation in 5 months old control iPSC-derived astrocytes. A control IgG antibody or an antibody directed against the C-terminal domain of LRP10 were used for immunoprecipitation. Flowthrough = FT. **D**. Representative confocal images of LRP10, NDFIP1, and CD63 in 4 months old control iPSC-derived astrocytes. Zoomed-in images are shown at the bottom of each panel. Yellow arrowheads show LRP10 and NDFIP1 overlap. White arrowheads show LRP10, NDFIP1 and CD63 overlap. Zoomed images scale bar = 5 µm. **E**. Representative western blot of cell lysates and conditioned media from HEK-293T cells overexpressing combinations of LRP10, NDFIP1, and α-synuclein. **F**. Quantification of secreted α-synuclein normalized to intracellular vinculin from E. n = 4 biological replicates. **G**. Quantification of intracellular α-synuclein normalized to vinculin from E. n = 4 biological replicates. All data are expressed as mean ± SD with individual data points shown. Data were analyzed by One-way ANOVA with Tukey’s multiple comparisons test. *P ≤ 0.05, **P ≤ 0.01, ***P ≤ 0.001, ****P ≤ 0.0001.

To validate the NDFIP1-LRP10 interaction under more physiological conditions, we performed LRP10 immunoprecipitation in cell lysates from control and LRP10-KO iPSC-derived astrocytes (Fig. 3B). We observed that NDFIP1 co-immunoprecipitated with LRP10 in control and not in LRP10-KO astrocytes (Fig. 3B). Additionally, we performed an immunoprecipitation with extracts from control iPSC-derived astrocytes using a control IgG or KO-validated anti-LRP10 antibody (Grochowska et al., 2021), which also verified the NDFIP1:LRP10 interaction (Fig. 3C).

To study the subcellular distribution of endogenously expressed LRP10 and NDFIP1, we next carried out immunocytochemistry of LRP10 and NDFIP1 relative to CD63 in control astrocytes (Fig. 3D). CD63 is a member of the tetraspanin protein family that resides in late endosomes, lysosomes, secretory vesicles, and at the plasma membrane, and moves among these compartments (Pols and Klumperman, 2009). LRP10 and NDFIP1 expression strongly overlapped at the perinuclear region or in vesicles (Fig. 3D). Furthermore, several LRP10 and NDFIP1-positive vesicles were also CD63-positive (Fig. 3D). To conclude, these data show a high degree of overlap between LRP10 and NDFIP1 in vesicular structures, in support of the NDFIP1:LRP10 interaction.

Since both LRP10 and NDFIP1 are associated with the secretory pathway (Carreras Mascaro et al., 2023; Putz et al., 2012; Putz et al., 2008) and α-synuclein processing (Borland et al., 2022; Carreras Mascaro et al., 2023; Liu et al., 2020), we investigated whether these pathways might converge. LRP10, NDFIP1, and α-synuclein were overexpressed in HEK-293T cells and intra- and extracellular α-synuclein levels were analyzed 48 h after transfection (Fig. 3E). We observed a significant 7-fold increase in α-synuclein secretion in cells overexpressing both LRP10 and α-synuclein in comparison to cells overexpressing α-synuclein alone, which is in line with our previous findings (Carreras Mascaro et al., 2023). Strikingly, overexpressing NDFIP1 and α-synuclein, and a combination of LRP10, NDFIP, and α-synuclein resulted in a comparable increase in extracellular α-synuclein levels (Fig. 3F). Additionally, LRP10-OE led to a decrease in intracellular α-synuclein levels (1,4-fold), which was further reduced via co-overexpression of NDFIP1 (3-fold, Fig. 3G). Interestingly, we also observed that NDFIP1-OE lead to changes in LRP10 levels in the triple-overexpressing cells (Supplementary Fig. 3A), and both LRP10-OE and α-synuclein-OE lead to changes in NDFIP1 levels (Supplementary Fig. 3B). Together, these results suggest that NDFIP1 and LRP10 are both implicated in regulating α-synuclein levels and secretion possibly via converging mechanisms.

### The LRP10 interactome in LRP10 variant-carrier iPSC-derived astrocytes

Previous work demonstrated that the *LRP10*:c.1424+5G>A variant (LRP10-Splice, Table 1) identified in an individual with DLB (Quadri et al., 2018), which encodes a truncated LRP10 protein lacking most of its functional domains, has a dominant negative effect on LRP10-mediated α-synuclein secretion (Carreras Mascaro et al., 2023). To determine whether this dominant negative effect might be accompanied by a dysregulation of the LRP10 interactome we performed LRP10 co-immunoprecipitations followed by MS on cell extracts from two iPSC-derived astrocytes clones from a donor carrying the LRP10-Splice variant. We identified 111 proteins interacting with LRP10 in LRP10-Splice astrocytes (Supplementary Table 2B, C), among which 80 were also detected in LRP10-WT astrocytes (Fig 4A). Interestingly, NDFIP1 was not found as an LRP10 interactor in either of the two LRP10-Splice astrocyte clones (Supplementary Table 2B). Seventy-one GOBP terms grouped into 9 summarized terms were significantly enriched (*BHp*-value < 0.05) in the LRP10-Splice interactome (Fig 4B, Supplementary Table 5). From these, 50 GOBP terms and 4 summarized terms were shared between both LRP10-WT and LRP10-Splice interactomes (Fig 4B, 4C). These terms were cytoplasmic translation, actin filament capping, negative regulation of protein-containing complex disassembly, and actin filament-based movement. On the other hand, 5 terms were only enriched in the proteins detected in LRP10-Splice astrocytes, including negative regulation of cytoskeleton organization, muscle development, non-membrane-bounded organelle assembly, wound healing, and regulation of body fluid levels (Fig 4C). Taken together, these data point towards possible differences in biological processes between LRP10-Splice carriers and controls, suggesting potential relevance in LBD mechanisms.

**Figure 4:**
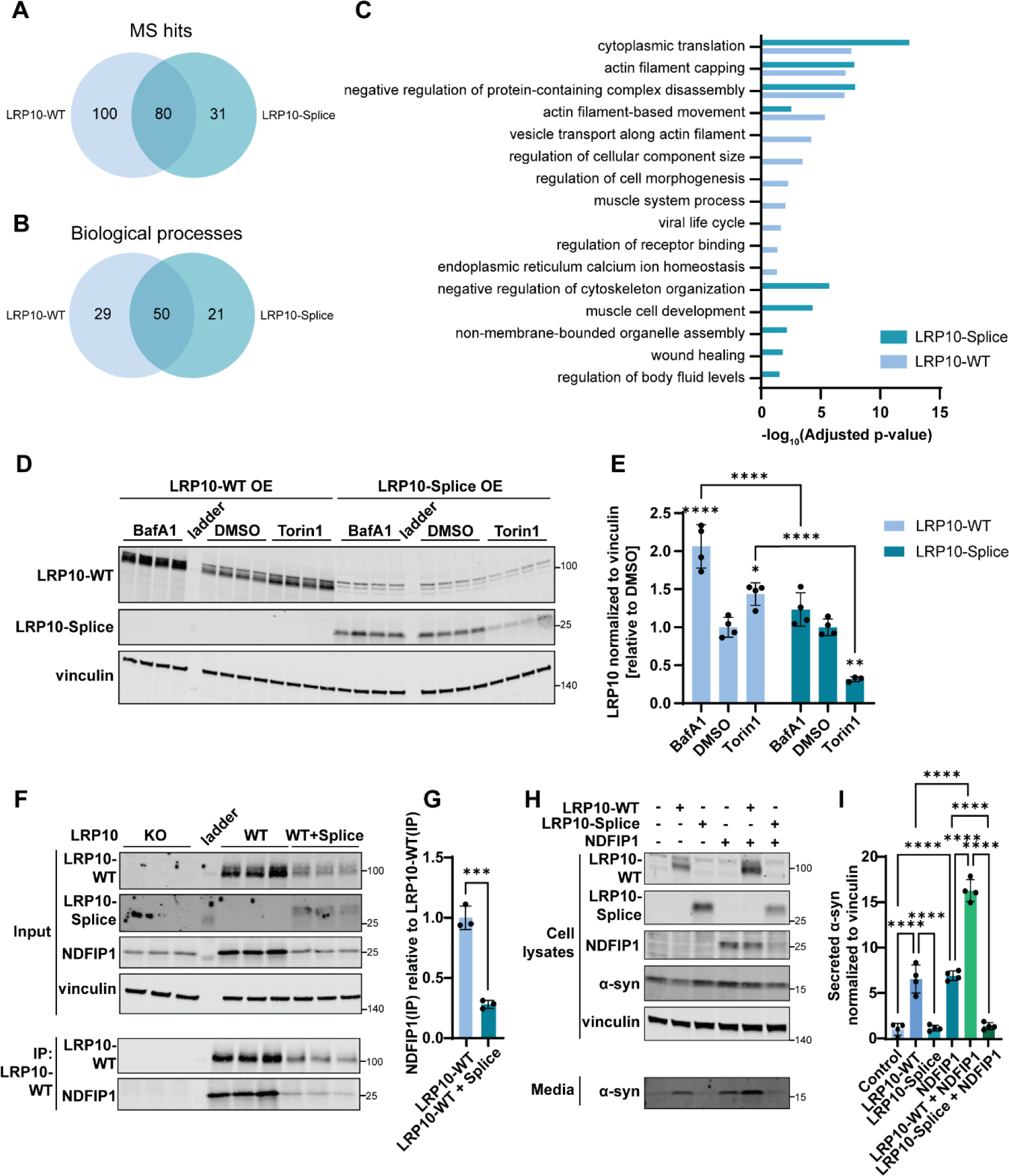
LRP10 interactome MS in patient-derived astrocyte lines. **A**. Venn diagram showing the overlap between the proteins identified in the whole-cell extracts of LRP10-WT and LRP10-Splice 4 months old iPSC-derived astrocytes. **B**. Venn diagram showing the overlap between the biological processes significantly enriched in the whole-cell extracts of LRP10-WT and LRP10-Splice iPSC-derived astrocytes **C**. Bar chart showing the summarized statistically significant GOBP identified in LRP10-WT and LRP10-Splice iPSC-derived astrocytes LRP10 co-immunoprecipitations/MS. For each summarized category, the lowest BHp-value among the terms in that category was used for plotting. **D**. Western blot of HEK-293T cells with a LRP10-KO background overexpressing LRP10-WT or LRP10-Splice and treated for 4h with BafA1, DMSO, or Torin1. **E**. Western blot quantification of LRP10 normalized to vinculin relative to DMSO for each transfection type from D. n = 3 or 4 biological replicates. **F**. Western blot of LRP10-WT and NDFIP1 co-immunoprecipitation in LRP10-KO, LRP10-WT-OE, and LRP10-WT+LRP10-Splice-OE HEK-293T cells. An antibody directed against the C-terminal domain of LRP10 that does not bind to LRP10-Splice was used for immunoprecipitation. **G**. Quantification of NDFIP1 that co-immunoprecipitated with LRP10-WT normalized to the pulled LRP10-WT (IP). n = 3 biological replicates. **H**. Representative western blot of cell lysates and conditioned media from HEK-293T cells overexpressing α-synuclein and combinations of LRP10-WT, LRP10-Splice, and NDFIP1. **I**. Quantification of secreted α-synuclein normalized to intracellular vinculin from H. n = 4 biological replicates. All data are expressed as mean ± SD with individual data points shown. Data from E were analyzed by Two-way ANOVA with Tukey’s multiple comparisons test. Data from G were analyzed by student’s T test. Data from I were analyzed One-way ANOVA with Sidak’s multiple comparisons test. *P ≤ 0.05, **P ≤ 0.01, ***P ≤ 0.001, ****P ≤ 0.0001.

### LRP10-Splice is aberrantly processed by the autophagy pathway and interferes with LRP10-WT and NDFIP1-mediated α-synuclein secretion

Considering the evidence that LRP10-WT plays an active role in autophagy and is degraded via the autophagy pathway (Fig. 2), we sought to investigate the effect of autophagy-modifying drugs on LRP10-Splice processing. We overexpressed LRP10-WT or LRP10-Splice in HEK-293T cells, treated them with BafA1 or Torin1 for 4h, and analyzed LRP10 protein expression levels (Fig. 4D). We observed significantly increased LRP10-WT levels after BafA1 (2-fold) and Torin1 (1.5-fold) treatments (Fig. 4E). However, LRP10-Splice levels did not change after BafA1 treatment and showed a 4-fold reduction after Torin1 treatment (Fig. 4E). Therefore, these results indicate that LRP10-Splice is aberrantly processed by the autophagy pathway.

LRP10-Splice exerts a dominant negative effect over LRP10-mediated α-synuclein secretion (Carreras Mascaro et al., 2023). Based on data showing that NDFIP1 is involved in regulating α-synuclein secretion (Fig. 3E-G) and that the NDFIP1:LRP10 interaction was not detected in LRP10-Splice variant-carrying astrocytes (Supplementary Table 2B), we investigated the effect of LRP10-Splice expression on the LRP10-WT:NDFIP1 interaction and NDFIP1-mediated α-synuclein secretion. First, we overexpressed NDFIP1 together with LRP10-WT or with a combination of LRP10-WT and LRP10-Splice cDNA (Carreras Mascaro et al., 2023) in LRP10-KO HEK-293T cells. LRP10-WT was immunoprecipitated from cell extracts using an antibody directed against the C-terminal domain of LRP10 that does not bind to LRP10-Splice, which lacks the C-terminal domain. Western blotting revealed that NDFIP1 co-immunoprecipitates with LRP10-WT in both conditions and not in the LRP10-KO control (Fig. 4F). However, a significantly lower amount of NDFIP1 co-immunoprecipitates with LRP10-WT when LRP10-Splice is present (4-fold, Fig. 4G). Together, these data provide evidence that LRP10-Splice interferes with the LRP10-WT:NDFIP1 interaction.

Next, we overexpressed α-synuclein with different combinations of LRP10-WT, LRP10-Splice, and NDFIP1 in LRP10-KO cells, and studied α-synuclein secretion 48 h after transfection via western blot (Fig. 4H). Overexpression of LRP10-WT or NDFIP1 alone induced a 6-fold increase in α-synuclein secretion (Fig. 4I). Strikingly, secreted α-synuclein levels increased 16-fold when LRP10 and NDFIP1 were overexpressed together (Fig. 4I). In contrast, not only was LRP10-Splice unable to induce an increase in α-synuclein secretion, but it also blocked the stimulatory effect of NDFIP1 overexpression on α-synuclein secretion (Fig. 4I). Taken together, these results indicate that LRP10-Splice interferes with the LRP10-WT:NDFIP1 interaction and inhibits LRP10-WT and NDFIP1-mediated α-synuclein secretion.

### LRP10 interactome in LRP10-Splice conditioned media

We previously showed that iPSC-derived astrocytes originating from a DLB patient carrying the *LRP10*:c.1424+5G>A (LRP10-Splice) variant secreted an aberrant high molecular weight species of LRP10 of unknown nature (Carreras Mascaro et al., 2023). Even though LRP10-WT is secreted via EVs, the patient-associated LRP10 species was mostly present in the EV-free media fraction (Carreras Mascaro et al., 2023). To further investigate the composition and protein interactors of this secreted EV-free LRP10 protein species, we fractionated conditioned media from LRP10-Splice iPSC-derived astrocytes via sequential ultracentrifugation and filtration according to published protocols (Carreras Mascaro et al., 2023) (Fig. 5A). Western blot inspection of cell lysates and the EV-free media fraction from control astrocytes and astrocytes derived from the DLB patient carrying the LRP10-Splice variant revealed the presence of the aberrant patient-specific LRP10 species (Fig. 5B). Next, we performed LRP10 immunoprecipitation and MS. We detected a total of 11 distinct peptides that mapped to the LRP10-WT sequence and no peptide mapping on the 33 amino acids sequence unique to the predicted product from LRP10-Splice (Fig 5C). This confirmed that the aberrant high molecular weight species observed in the EV-free media fraction is at least partially comprised of wild-type LRP10, even though control cells do not secrete LRP10 in the EVs-free fraction at detectable levels.

**Figure 5:**
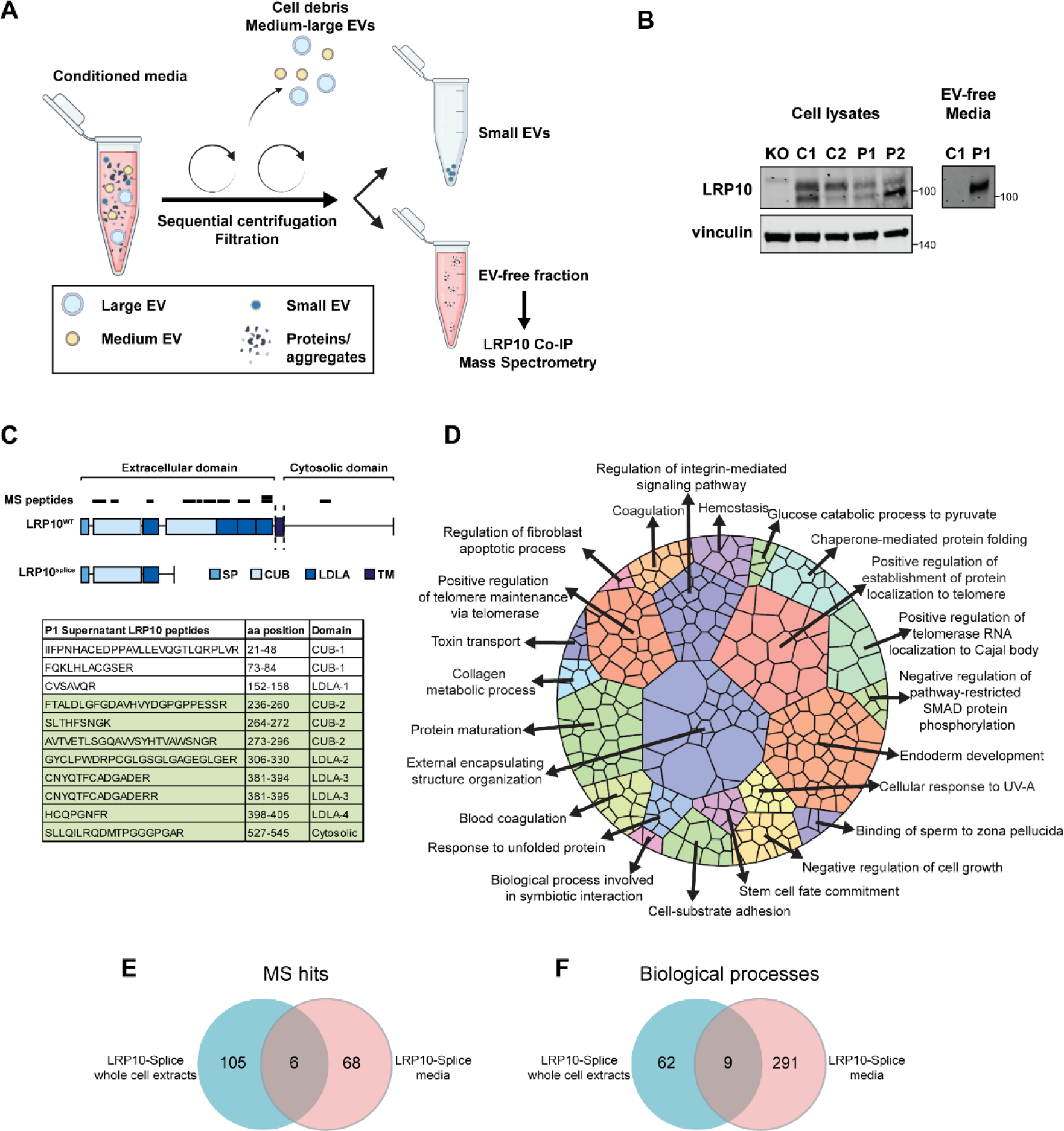
Secreted LRP10-Splice interactome MS. **A**. Schematic representation of the protocol used to fraction conditioned media to obtain the EV-free fraction. **B**. Western blot showing LRP10 in cell lysates and EV-free media fraction from two control and two clones from the LRP10-Splice carrier iPSC-derived astrocytes lines. **C**. Mapping of the peptides identified by MS on the sequences of LRP10-WT and LRP10-Splice, and a table indicating the peptides and the LRP10 domains they align to. The sequences that only map to LRP10-WT and not to LRP10-Splice are highlighted in green. **D**. Voronoi plot showing the 300 statistically significant GOBP enriched in the LRP10-Splice astrocyte conditioned media, grouped into the 23 summary terms. **E**. Venn diagram showing the overlap between the proteins identified in whole-cell extracts and EV-free media fraction in LRP10-Splice iPSC-derived astrocytes. **F**. Venn diagram showing the overlap between the significantly enriched biological processes in whole-cell extracts and EV-free media fraction in LRP10-Splice iPSC-derived astrocytes.

From the MS analysis, we identified 74 proteins in the LRP10-Splice EV-free media (Supplementary Table 2C). The LRP10 interactome in the EV-free media fraction was significantly enriched (*BHp*-value < 0.05) for 300 GOBP terms belonging to 23 summarized terms (Fig. 5D, Supplementary Table 6), e.g. external encapsulating structure organization, chaperone-mediated protein folding, protein maturation, and cell-substrate adhesion. Only six of the proteins identified in the LRP10-Splice astrocytes media were found also in the LRP10-Splice whole-cell extracts (Fig. 5E, Supplementary Table 2C). Nine terms were enriched in both whole-cell extracts and media (Fig. 5F). These terms included cell junction assembly, homotypic cell-cell adhesion, osteoblast differentiation, peptide cross-linking, plasma membrane repair, regulation of body fluid levels, regulation of cell growth, regulation of cell-substrate adhesion, and wound healing while none of the summarized GOBP terms was shared. These results implicate that the aberrantly secreted LRP10-Splice species contains LRP10-WT and it is associated with proteins that might be relevant to understand LRP10 cell-to-cell transmission and downstream pathological pathways.

## Discussion

An increasing amount of evidence has positioned LRP10 as a protein of interest in LBDs and other neurodegenerative diseases, but its physiological function as well as its role in neurodegeneration remain incompletely characterized. In the current study, we aimed at expanding our understanding of the biology of LRP10 by mapping its interactome in an overexpression model and iPSC-derived astrocytes expressing endogenous LRP10. In addition to identifying biological processes associated with the LRP10 interactome, we characterized its role in p62-mediated autophagy and NDFIP1-mediated α-synuclein processing. Furthermore, we identified LRP10 interactors in cell lysates and conditioned media of iPSC-derived astrocytes originating from a *LRP10*:c.1424+5G>A (LRP10-Splice) variant-carrying DLB patient, and showed that this variant affects LRP10 processing via the autophagy pathway and interferes with the LRP10:NDFIP1 interplay.

Some of the LRP10-interactors we report in this work have been previously described. LRP10 was found to interact with CISD2, EBP, MGST3 by two-hybrid screening (Luck et al., 2020). Moreover, several studies using affinity-capture followed by MS listed as LRP10-interactors GGA2 (Sahgal et al., 2019), CDC42 (Bagci et al., 2020), SPINT2, HEATR3, NDFIP2, PCDHA8, SYT11, TMEM59 (Huttlin et al., 2021; Huttlin et al., 2017; Huttlin et al., 2015), and WWP1 (Huttlin et al., 2021; Huttlin et al., 2017; Huttlin et al., 2015; Nielsen et al., 2019). The fact that the majority of the LRP10-interactors we present here have not been identified in previous studies, and several previously reported interactors were not identified in this study, is probably due to the difficulties in mapping the human proteome. The high number of proteins, isoforms, and modifications combined with differences in expression that also depend on the specific biological systems, and the prevalence of transient interactions are just a few biological reasons that hinder a complete protein interactome mapping (Huttlin et al., 2021). Moreover, the use of different experimental approaches, models, protocols, and the presence of tags that are pivotal for the purification, but can mask the sites needed for biologically relevant interactions with other proteins can bias the final mapping. To tackle some of these limitations, in the present work, we used overexpressed untagged full-length LRP10 cDNA or endogenous LRP10 in a biologically relevant system, and a KO-validated antibody directed against the N-terminal domain of LRP10 (Grochowska et al., 2021). One argument in favor of using untagged LRP10 relies on LRP10 structure. LRP10 is a single-pass transmembrane protein (Pohlkamp et al., 2017), which presents two motifs consisting of a cluster of acidic amino-acids followed by two leucine residues (DXXLL) close to its C-terminal end that have been shown to be essential for LRP10 localization (Boucher et al., 2008; Doray et al., 2008; Quadri et al., 2018). We speculate that the presence of a protein tag in the vicinity of these motifs might affect the binding of LRP10 to some of its interactors. Therefore, our approach to study the LRP10 interactome without protein tags at endogenous expression levels in iPSC-derived astrocytes, which is a physiologically relevant cell model, address some of the experimental confounding factors and supports the detection of additional physiological LRP10 protein interactors. LRP10 function has been associated with the intracellular vesicle transport pathway due to its interaction with the retromer complex and clathrin adaptors and its intracellular distribution pattern (Boucher et al., 2008; Brodeur et al., 2009; Daly et al., 2023; Doray et al., 2008; Grochowska et al., 2021; Quadri et al., 2018; Steinberg et al., 2013). Of note, LRP10 trafficking has also been found to be regulated by the Ca^2+^ sensor nucleobindin 1 (Brodeur et al., 2009). Furthermore, drugs targeting the integrity of the ER-Golgi, ER and proteasomal stress, and autophagic function have been shown to influence LRP10 levels, activity, and/or localization (Boucher et al., 2008; Carreras Mascaro et al., 2023). Our MS analyses revealed several LRP10-interacting proteins related to the previously described pathways included in the following summarized gene ontology terms: “establishment of protein localization to organelle”, “response to endoplasmic reticulum stress”, “proteasome-mediated ubiquitin-dependent protein catabolic process”, “selective autophagy” (Fig. 1A, Supplementary Table 3), “vesicle transport along actin filament”, and “endoplasmic reticulum calcium ion homeostasis” (Fig.1C, Supplementary Table 4). Thus, our data are in line with previous studies on LRP10 protein function, and also provides additional information to build more complex and detailed LRP10-associated biological processes.

In a recent study, we reported that LRP10 expression levels and secretion depend on autophagic function (Carreras Mascaro et al., 2023). Specifically, BafA1 was shown to induce the accumulation of LRP10 in vesicular structures and Torin1 was shown to decrease LRP10 levels, suggesting autophagic degradation (Carreras Mascaro et al., 2023). Immunocytochemistry of LRP10 and LC3B in iPSC-derived astrocytes showed an increase in their colocalization after BafA1 or BafA1+Torin1 treatments, indicating accumulation of both proteins, and a decrease in their colocalization after Torin1 treatment, suggesting increased degradation (Fig. 2C-E). Interestingly, LRP10 and LC3B-positive structures seem to be in close proximity with each other rather than perfectly overlapping, which is especially visible after BafA1+Torin1 treatment, suggesting the interaction of two types of vesicular structures (Fig. 2C, Zoom). This is reminiscent of the close apposition of endosomes (LRP10-positive) and autophagosomes (LC3B-positive) forming amphisomes (Zhao et al., 2021b). Localization of LRP10 in the endosomal pathway has been extensively reported, and is also in line with the observed colocalization between LRP10 and CD63 (Fig. 3D). More detailed analyses are required to further delineate LRP10 localization in these structures. We also uncovered that LRP10 physically interacts with and regulates expression levels of the autophagy receptor SQSTM1/p62 (Fig. 2A, 2B), supporting a role for LRP10 in the autophagy pathway. P62 is involved in the formation of autophagosomes, serves as a receptor for the delivery of ubiquitinated protein cargos to lysosomes or the ubiquitin proteasome system, as well as the formation of stable intracellular inclusions (Trejo-Lopez et al., 2020). Whether and how LRP10 and p62 affect each other in these processes remains to be determined. Interestingly, our results from LRP10 knock-out and overexpressing cells suggest that LRP10 promotes autophagic degradation (Fig. 2F-K). Defects in the autophagy-lysosomal pathway are a recurrent theme in LBDs (Fellner et al., 2021). For instance, SYNJ1/Synaptojanin1 is required for macroautophagy, a role that is inhibited by the PD-causing mutation R258Q (Quadri et al., 2013; Vanhauwaert et al., 2017). Furthermore, recent studies demonstrate that loss of DNAJC6/auxilin, another gene with variants causing juvenile/early onset PD (Edvardson et al., 2012), leads to increased pre-synaptic macroautophagy (Vidyadhara et al., 2023). Moreover, monogenic PD-associated genes *PINK1*, encoding the PTEN-induced serine/threonine kinase 1, and *PRKN*, encoding an E3 ubiquitin ligase Parkin, regulate mitochondrial quality control via a selective autophagy pathway termed mitophagy (Wang et al., 2022). Importantly, LBD-associated genetic defects in *GBA* and *LRRK2* have been shown to induce severe defects in autophagy and lysosomal degradation, leading to accumulation of toxic material, and ultimately to neuronal death (Bae et al., 2015; Du et al., 2015; Henry et al., 2015; Kania et al., 2023; Pang et al., 2022; Schapansky et al., 2018). Taken together, our data implicates LRP10 in the autophagy pathway, strengthening the connection to LBDs disease-associated mechanisms previously reported for LBD-causing genes. Future research should delineate the role of LRP10, the effect of LRP10 disease-associated variants, and the potential interactions between LRP10 and the products of other LBD-associated genes in this pathway.

We identified NDFIP1 as a novel LRP10 interactor in both HEK-293T cells and iPSC-derived astrocytes (Fig. 1, 2), and demonstrated a high level of overlap of LRP10 and NDFIP1 in vesicles (Fig. 3D). Previous reports suggest a possible association of NDFIP1 with LBD pathogenesis. NDFIP1 is a HECT-type ubiquitin E3 ligase adaptor and activator known to be involved in α-synuclein degradation (Borland et al., 2022) and it was suggested to have a neuroprotective role in a cellular model of PD (Liu et al., 2020). Importantly, NDFIP1 was shown to be elevated in substantia nigra of PD patients, and it was ectopically localized in astrocytes (Howitt et al., 2014). Interestingly, both LRP10 (Grochowska et al., 2021) and NDFIP1 (Howitt et al., 2014) were shown to localize to α-synuclein-positive inclusions in dopaminergic neurons in PD brains. Moreover, both LRP10 and NDFIP1 are associated with EVs and the secretion of α-synuclein (Carreras Mascaro et al., 2023; Howitt et al., 2021; Putz et al., 2012; Putz et al., 2008). Here, we show that both NDFIP1 and LRP10 overexpression can increase extracellular α-synuclein levels to a similar extent (Fig. 3E, 3F, 4F, 4G). Furthermore, NDFIP1 OE amplified the effect of LRP10 OE decreasing intracellular α-synuclein levels (Fig. 3E, 3G). Interestingly, LRP10 was previously shown to induce α-synuclein secretion via a proteasome-dependent pathway (Carreras Mascaro et al., 2023). Given that NDFIP1 is well known for its role as a ubiquitin-proteasome adaptor (Low et al., 2015; Putz et al., 2012), it is possible that both LRP10 and NDFIP1 regulate α-synuclein secretion via a proteasome-mediated mechanism, and further research is warranted to further elucidate it.

The LBD-associated LRP10-Splice variant was suggested to have a dominant negative effect over LRP10-WT and to interfere with LRP10-mediated α-synuclein secretion (Carreras Mascaro et al., 2023). Here, we explored the effect of this variant on the LRP10 interactome and associated biological processes (Fig. 4). A comparison at the biological process level was performed, as the proteomics approach we employed is not quantitative and direct comparison of the proteomics hits across conditions is not possible. In addition to shared GO terms between LRP10-WT and LRP10-Splice, also differences could be identified. For instance, ontologies such as “vesicle transport along actin filament”, “regulation of receptor binding”, and “endoplasmic reticulum calcium ion homeostasis”, which have been shown to be relevant for LRP10 function in other studies (Boucher et al., 2008; Brodeur et al., 2009; Carreras Mascaro et al., 2023; Doray et al., 2008; Grochowska et al., 2021; Quadri et al., 2018; Steinberg et al., 2013), were only present in control cells but not in patient-derived cells (Fig. 4). These findings point to possible defects in these pathways in patient-derived cells, which remains to be experimentally validated in future research. Nevertheless, the current work provides initial clues of the effect of the LRP10-Splice variant on the LRP10 interactome.

Next, to further characterize possible defects associated with the LRP10-Splice variant, we overexpressed the patient’s cDNA in HEK-293T cells. Importantly, a different LRP10 variant in the same genomic position identified in three PD patients, *LRP10*:c.1424+5delG (Quadri et al., 2018), results in the same truncated cDNA species. Here, we investigated the effect of autophagy drugs on LRP10-Splice processing and we observed that, unlike LRP10-WT, LRP10-Splice levels remained unchanged to BafA1 treatment (Fig. 4D and 4E), suggesting that this variant is not degraded in lysosomes. Interestingly, LRP10-Splice levels were significantly reduced upon Torin1 treatment, indicating that promoting autophagic function leads to the disposal of this truncated protein in a lysosome-independent manner. Torin1 is an mTOR kinase inhibitor, and mTOR inhibition has been shown to activate overall protein degradation not only by autophagy, but by the ubiquitin-proteasome system as well, possibly explaining the LRP10-Splice reduction after Torin1 treatment (Zhao et al., 2015). Since NDFIP1 was not detected as a LRP10 interactor in LRP10-Splice astrocytes via MS (Supplementary Table 2B, 1C), we studied the effect of LRP10-Splice OE on the LRP10:NDFIP1 interaction. We found that LRP10-Splice reduces the amount of NDFIP1 that co-immunoprecipitated with LRP10-WT, possibly by competition between LRP10-Splice and NDFIP1 for binding to LRP10-WT (Fig. 4F, 4G). Furthermore, overexpression experiments in HEK-293T cells demonstrate that LRP10-Splice blocks NDFIP1-mediated α-synuclein secretion (Fig. 4H, 4I). Taken together, we show that LRP10 modulates the secretion of α-synuclein via interaction with NDFIP1, provide further evidence for the dominant negative effect of LRP10-Splice (Carreras Mascaro et al., 2023) on the LRP10:NDFIP1 interaction and α-synuclein secretion, and aberrant LRP10-Splice processing via the autophagy pathway. Altogether, these findings suggest defects in the autophagic pathway and α-synuclein processing as possible disease mechanisms in patients carrying LRP10 mutations in position c.1424+5.

Finally, we show that the previously reported high molecular weight LRP10 species only detected in the EV-free fraction of the conditioned media of LRP10-Splice-expressing astrocytes (Carreras Mascaro et al., 2023) is constituted in part by full-length wild-type LRP10 (Fig. 5). Moreover, the interactome of this patient-derived extracellular LRP10 species only minimally overlaps with the interactors identified in whole-cell extracts from the same cells. Interestingly, several of the proteins interacting with this patient-associated LRP10 species function in protein folding after the detection of unfolded proteins, possibly indicating that this LRP10 form presents an aberrant structure. Future research is required to dissect the possible impact of the patient-associated secreted LRP10 species.

In conclusion, we have mapped the LRP10 interactome in LRP10-overexpressing cells and in iPSC-derived astrocytes with high endogenous levels of LRP10 (Grochowska et al., 2021). We used these datasets to uncover and confirm the involvement of LRP10 in autophagy, potentially by its interaction with p62, and further characterized the LRP10-mediated regulation of α-synuclein levels and secretion via the newly identified interaction between LRP10 and NDFIP1. We also nominate a series of patient-specific biological processes both intracellularly and in the EV-free media fraction where the patient-specific LRP10 species might be involved. Furthermore, we show that autophagy-mediated processing of LRP10 is affected by the presence of the LRP10-Splice variant, and that LRP10-Splice expression interferes with the LRP10:NDFIP1 interaction. This study provides new insights into LRP10 function and its role in neurodegeneration and provides identifies promising target pathways for potential therapeutic disease-modifying interventions.

## Supporting information

SupplementaryTable_1_Maxquant_files

SupplementaryTable_2_Overview_hits

SupplementaryTable_3_HEK_GOBP

SupplementaryTable_4_Astro-WT_GOBP

SupplementaryTable_5_Astro-Splice_GOBP

SupplementaryTable_6_Astro-Splice_media_GOBP

## Data availability

MS raw data were submitted as supplemental tables to the ProteomeXchange Consortium via the PRIDE partner repository (Perez-Riverol et al., 2019) with the data identifier PXD045871. For reviewing purposes, the following credentials can be used:

Username

Password: HQSXX7ah

## Acknowledgments

We thank Martyna M. Grochowska for providing the LRP10-KO and NPC cell lines, sharing the iPSC-derived astrocytes differentiation protocol, and providing insight that greatly assisted the research in this manuscript. We would also like to show our gratitude to the Erasmus MC iPS Core Facility for providing the iPSC lines.

## Author contributions

A.C.M. and W.M. designed the experiments. A.C.M., V.Boumeester., G.J.B., D.H.W.D., and J.A.A.D. performed the experiments. A.C.M., F.F., D.H.W.D., and J.A.A.D. performed data analysis. A.C.M., F.F., and W.M interpreted the data. L.J.M.V. and F.J.d.J. provided the patient biopsy. A.C.M. and F.F. wrote the initial draft of the manuscript. V.Bonifati and W.M. conceived the study and supervised the experiments. All authors critically read the manuscript and approved the final version.

## References

Attems, J., et al., 2021. Neuropathological consensus criteria for the evaluation of Lewy pathology in post-mortem brains: a multi-centre study. Acta Neuropathologica. 141, 159–172.

Bae, E. J., et al., 2015. Loss of glucocerebrosidase 1 activity causes lysosomal dysfunction and α-synuclein aggregation. Exp Mol Med. 47, e153.

Bagci, H., et al., 2020. Mapping the proximity interaction network of the Rho-family GTPases reveals signalling pathways and regulatory mechanisms. Nat Cell Biol. 22, 120–134.

Barbar, L., et al., 2020. CD49f Is a Novel Marker of Functional and Reactive Human iPSC-Derived Astrocytes. Neuron. 107, 436–453.e12.

Behl, T., et al., 2022. Exploring the Role of Ubiquitin–Proteasome System in Parkinson’s Disease. Molecular Neurobiology. 59, 4257–4273.

Borland, H., et al., 2022. α-synuclein buildup is alleviated via ESCRT-dependent endosomal degradation brought about by p38MAPK inhibition in cells expressing p25α. J Biol Chem. 298, 102531.

Boucher, R., et al., 2008. Intracellular trafficking of LRP9 is dependent on two acidic cluster/dileucine motifs. Histochem Cell Biol. 130, 315–27.

Brodeur, J., et al., 2009. Calnuc binds to LRP9 and affects its endosomal sorting. Traffic. 10, 1098–114.

Brodeur, J., et al., 2012. LDLR-related protein 10 (LRP10) regulates amyloid precursor protein (APP) trafficking and processing: evidence for a role in Alzheimer’s disease. Mol Neurodegener. 7, 31.

Burré, J., et al., 2018. Cell Biology and Pathophysiology of α-Synuclein. Cold Spring Harbor Perspectives in Medicine. 8.

Carreras Mascaro, A., et al., 2023. LRP10 as a novel α-synuclein regulator in Lewy body diseases. bioRxiv. 2023.05.12.540510.

Chen, Y., et al., 2019. LRP10 in autosomal-dominant Parkinson’s disease. Mov Disord. 34, 912–916.

Cox, J., et al., 2011. Andromeda: A Peptide Search Engine Integrated into the MaxQuant Environment. Journal of Proteome Research. 10, 1794–1805.

Daida, K., et al., 2019. Mutation analysis of LRP10 in Japanese patients with familial Parkinson’s disease, progressive supranuclear palsy, and frontotemporal dementia. Neurobiol Aging. 84, 235.e11–235.e16.

Daly, J. L., et al., 2023. Multi-omic approach characterises the neuroprotective role of retromer in regulating lysosomal health. Nature Communications. 14, 3086.

Doray, B., et al., 2008. Identification of acidic dileucine signals in LRP9 that interact with both GGAs and AP-1/AP-2. Traffic. 9, 1551–62.

Du, T.-T., et al., 2015. GBA deficiency promotes SNCA/α-synuclein accumulation through autophagic inhibition by inactivated PPP2A. Autophagy. 11, 1803–1820.

Edvardson, S., et al., 2012. A deleterious mutation in DNAJC6 encoding the neuronal-specific clathrin-uncoating co-chaperone auxilin, is associated with juvenile parkinsonism. PLoS One. 7, e36458.

Fellner, L., et al., Autophagy in α-Synucleinopathies—An Overstrained System. Cells, Vol. 10, 2021.

Gagliardi, M., et al., 2021. Analysis of the LRP10 gene in patients with Parkinson’s disease and dementia with Lewy bodies from Southern Italy. Neurol Sci. 42, 305–308.

Goralski, T., et al., 2023. Spatial transcriptomics reveals molecular dysfunction associated with Lewy pathology. bioRxiv. 2023.05.17.541144.

Grochowska, M. M., et al., 2022. CRISPR/Cas9-mediated LRP10 Knockout in HuTu-80 and HEK 293T Cell Lines. Bio-protocol. 12, e4521.

Grochowska, M. M., et al., 2021. LRP10 interacts with SORL1 in the intracellular vesicle trafficking pathway in non-neuronal brain cells and localises to Lewy bodies in Parkinson’s disease and dementia with Lewy bodies. Acta Neuropathol. 142, 117–137.

Guo, L., et al., 2023. Sex specific molecular networks and key drivers of Alzheimer’s disease. Molecular Neurodegeneration. 18, 39.

Henry, A. G., et al., 2015. Pathogenic LRRK2 mutations, through increased kinase activity, produce enlarged lysosomes with reduced degradative capacity and increase ATP13A2 expression. Human Molecular Genetics. 24, 6013–6028.

Hou, X., et al., 2020. Autophagy in Parkinson’s Disease. Journal of Molecular Biology. 432, 2651–2672.

Howitt, J., et al., 2014. Increased Ndfip1 in the substantia nigra of Parkinsonian brains is associated with elevated iron levels. PLoS One. 9, e87119.

Howitt, J., et al., 2021. Exosomal transmission of α-synuclein initiates Parkinson’s disease-like pathology. bioRxiv. 2021.05.10.443522.

Huttlin, E. L., et al., 2021. Dual proteome-scale networks reveal cell-specific remodeling of the human interactome. Cell. 184, 3022–3040 e28.

Huttlin, E. L., et al., 2017. Architecture of the human interactome defines protein communities and disease networks. Nature. 545, 505–509.

Huttlin, E. L., et al., 2015. The BioPlex Network: A Systematic Exploration of the Human Interactome. Cell. 162, 425–440.

Kania, E., et al., 2023. LRRK2 phosphorylation status and kinase activity regulate (macro)autophagy in a Rab8a/Rab10-dependent manner. Cell Death & Disease. 14, 436.

Klionsky, D. J., et al., 2021. Guidelines for the use and interpretation of assays for monitoring autophagy (4th edition)1. Autophagy. 17, 1–382.

La Manno, G., et al., 2016. Molecular Diversity of Midbrain Development in Mouse, Human, and Stem Cells. Cell. 167, 566–580.e19.

Li, C., et al., 2021. Mutation analysis of LRP10 in a large Chinese familial Parkinson disease cohort. Neurobiol Aging. 99, 99.e1–99.e6.

Liao, T. W., et al., 2021. Role of LRP10 in Parkinson’s disease in a Taiwanese cohort. Parkinsonism Relat Disord. 89, 79–83.

Liu, X., et al., 2020. Ndfip1 Prevents Rotenone-Induced Neurotoxicity and Upregulation of α-Synuclein in SH-SY5Y Cells. Front Mol Neurosci. 13, 613404.

Low, L.-H., et al., 2015. Nedd4 Family Interacting Protein 1 (Ndfip1) Is Required for Ubiquitination and Nuclear Trafficking of BRCA1-associated ATM Activator 1 (BRAT1) during the DNA Damage Response*. Journal of Biological Chemistry. 290, 7141–7150.

Luck, K., et al., 2020. A reference map of the human binary protein interactome. Nature. 580, 402–408.

Ma, S., et al., 2019. SQSTM1/p62: A Potential Target for Neurodegenerative Disease. ACS Chemical Neuroscience. 10, 2094–2114.

Mandegar, M. A., et al., 2016. CRISPR Interference Efficiently Induces Specific and Reversible Gene Silencing in Human iPSCs. Cell Stem Cell. 18, 541–53.

Manini, A., et al., 2021. Screening of LRP10 mutations in Parkinson’s disease patients from Italy. Parkinsonism Relat Disord. 89, 17–21.

Neff, R. A., et al., 2021. Molecular subtyping of Alzheimer’s disease using RNA sequencing data reveals novel mechanisms and targets. Sci Adv. 7.

Neri, M., et al., 2021. Parkinson’s disease-dementia in trans LRP10 and GBA variants: Response to deep brain stimulation. Parkinsonism Relat Disord. 92, 72–75.

Ni, J., et al., 2021. Rare, pathogenic variants in LRP10 are associated with amyotrophic lateral sclerosis in patients from mainland China. Neurobiol Aging. 97, 145.e17–145.e22.

Nielsen, C. P., et al., 2019. USP9X Deubiquitylates DVL2 to Regulate WNT Pathway Specification. Cell Rep. 28, 1074–1089 e5.

Pang, S. Y.-Y., et al., 2022. LRRK2, GBA and their interaction in the regulation of autophagy: implications on therapeutics in Parkinson’s disease. Translational Neurodegeneration. 11, 5.

Perez-Riverol, Y., et al., 2019. The PRIDE database and related tools and resources in 2019: improving support for quantification data. Nucleic Acids Research. 47, D442–D450.

Periñán, M. T., et al., 2020. Analysis of p.Tyr307Asn variant in the LRP10 gene in Parkinson’s disease in southern Spain. Neurobiol Aging. 93, 142.e1–142.e3.

Pohlkamp, T., et al., 2017. Functional Roles of the Interaction of APP and Lipoprotein Receptors. Frontiers in Molecular Neuroscience. 10.

Pols, M. S., Klumperman, J., 2009. Trafficking and function of the tetraspanin CD63. Experimental Cell Research. 315, 1584–1592.

Putz, U., et al., 2012. The Tumor Suppressor PTEN Is Exported in Exosomes and Has Phosphatase Activity in Recipient Cells. Science Signaling. 5, ra70–ra70.

Putz, U., et al., 2008. Nedd4 family-interacting protein 1 (Ndfip1) is required for the exosomal secretion of Nedd4 family proteins. J Biol Chem. 283, 32621–7.

Quadri, M., et al., 2013. Mutation in the SYNJ1 Gene Associated with Autosomal Recessive, Early-Onset Parkinsonism. Human Mutation. 34, 1208–1215.

Quadri, M., et al., 2018. LRP10 genetic variants in familial Parkinson’s disease and dementia with Lewy bodies: a genome-wide linkage and sequencing study. Lancet Neurol. 17, 597–608.

Reinhardt, P., et al., 2013. Derivation and expansion using only small molecules of human neural progenitors for neurodegenerative disease modeling. PloS one. 8, e59252–e59252.

Sahgal, P., et al., 2019. GGA2 and RAB13 promote activity-dependent β1-integrin recycling. J Cell Sci. 132.

Sap, K. A., et al., 2017. Quantitative Proteomics Reveals Extensive Changes in the Ubiquitinome after Perturbation of the Proteasome by Targeted dsRNA-Mediated Subunit Knockdown in Drosophila. Journal of Proteome Research. 16, 2848–2862.

Sayols, S., rrvgo: a Bioconductor package for interpreting lists of Gene Ontology terms. microPublication Biology, 2023.

Schapansky, J., et al., 2018. Familial knockin mutation of LRRK2 causes lysosomal dysfunction and accumulation of endogenous insoluble α-synuclein in neurons. Neurobiol Dis. 111, 26–35.

Schwertman, P., et al., 2013. An immunoaffinity purification method for the proteomic analysis of ubiquitinated protein complexes. Analytical Biochemistry. 440, 227–236.

Song, N., et al., 2023. Genetic analysis of the LRP10 gene in Chinese patients with Parkinson’s disease. Neurol Sci. 44, 905–912.

Steinberg, F., et al., 2013. A global analysis of SNX27-retromer assembly and cargo specificity reveals a function in glucose and metal ion transport. Nat Cell Biol. 15, 461–71.

Sugiyama, T., et al., 2000. A novel low-density lipoprotein receptor-related protein mediating cellular uptake of apolipoprotein E-enriched beta-VLDL in vitro. Biochemistry. 39, 15817–25.

Tan, M. M. X., et al., 2019. Genetic analysis of Mendelian mutations in a large UK population-based Parkinson’s disease study. Brain. 142, 2828–2844.

Trejo-Lopez, J. A., et al., 2020. Generation and Characterization of Novel Monoclonal Antibodies Targeting p62/sequestosome-1 Across Human Neurodegenerative Diseases. Journal of Neuropathology & Experimental Neurology. 79, 407–418.

Vanhauwaert, R., et al., 2017. The SAC1 domain in synaptojanin is required for autophagosome maturation at presynaptic terminals. Embo J. 36, 1392–1411.

Vergouw, L. J. M., et al., 2020a. Clinical and Pathological Phenotypes of LRP10 Variant Carriers with Dementia. J Alzheimers Dis. 76, 1161–1170.

Vergouw, L. J. M., et al., 2020b. LRP10 variants in progressive supranuclear palsy. Neurobiol Aging. 94, 311.e5–311.e10.

Vergouw, L. J. M., et al., 2019. LRP10 variants in Parkinson’s disease and dementia with Lewy bodies in the South-West of the Netherlands. Parkinsonism Relat Disord. 65, 243–247.

Vidyadhara, D. J., et al., 2023. Dopamine transporter and synaptic vesicle sorting defects underlie auxilin-associated Parkinson’s disease. Cell Reports. 42, 112231.

Wang, J. Z., et al., 2007. A new method to measure the semantic similarity of GO terms. Bioinformatics. 23, 1274–81.

Wang, X.-L., et al., 2022. Mitophagy, a Form of Selective Autophagy, Plays an Essential Role in Mitochondrial Dynamics of Parkinson’s Disease. Cellular and Molecular Neurobiology. 42, 1321–1339.

Wu, T., et al., 2021. clusterProfiler 4.0: A universal enrichment tool for interpreting omics data. Innovation (Camb). 2, 100141.

Yu, G., 2020. Gene Ontology Semantic Similarity Analysis Using GOSemSim. Methods Mol Biol. 2117, 207–215.

Yu, G., et al., 2010. GOSemSim: an R package for measuring semantic similarity among GO terms and gene products. Bioinformatics. 26, 976–8.

Zhang, Y., et al., 2016. Purification and Characterization of Progenitor and Mature Human Astrocytes Reveals Transcriptional and Functional Differences with Mouse. Neuron. 89, 37–53.

Zhao, J., et al., 2015. mTOR inhibition activates overall protein degradation by the ubiquitin proteasome system as well as by autophagy. Proceedings of the National Academy of Sciences. 112, 15790–15797.

Zhao, Q., et al., 2021a. LRP10 Mutations May Correlate with Sporadic Parkinson’s Disease in China. Mol Neurobiol. 58, 1212–1216.

Zhao, Y., et al., 2020. The role of genetics in Parkinson’s disease: a large cohort study in Chinese mainland population. Brain. 143, 2220–2234.

Zhao, Y. G., et al., 2021b. Machinery, regulation and pathophysiological implications of autophagosome maturation. Nature Reviews Molecular Cell Biology. 22, 733–750.

