## Supplementary Tables Captions for "Interactome mapping reveals a role for LRP10 in autophagy and NDFIP1-mediated α-synuclein secretion"

**Supplementary Table 1:** Details of the proteomics hits detected in each experimental condition based on the MaxQuant 'proteingroups.txt' output files. Hits highlighted with the red font are the baits (LRP10) for the immune-precipitation and the hits highlighted in grey are annotated as ‘common contaminants’. **Sheet 1**: details of the first HEK293-T experiment. **Sheet 2**: details of the second HEK293-T experiment. **Sheet 3**: details of the LRP10-Splice iPSCs-derived astrocytes experiment. **Sheet 4**: details of the EV-free media conditioned by LRP10-WT iPSCs-derived astrocytes experiment.

**Supplementary Table 2:** Overview of the proteomics hits detected in each experimental condition and overlap with the others.

**Supplementary Table 3:** Results of the functional enrichment analysis of the LRP10-interactome in HEK293-T.

**Supplementary Table 4:** Results of the functional enrichment analysis of the LRP10-interactome in the LRP10-WT iPSCs-derived astrocytes.

**Supplementary Table 5:** Results of the functional enrichment analysis of the LRP10-interactome in the LRP10-Splice iPSCs-derived astrocytes.

**Supplementary Table 6:** Results of the functional enrichment analysis of the LRP10-interactome in the EV-free media conditioned by LRP10-Splice iPSCs-derived astrocytes.
